# BRD4 represses developmental and neuronal genes through interaction with PRC1.6

**DOI:** 10.64898/2026.01.31.702994

**Authors:** Fanny Boulet, Manthan Patel, Zeinab S. Zanjani, Nuria Andres-Sanchez, Anam Ijaz, Debosree Pal, Pankaj Dubey, Aoife Murray, Dean Nizetic, Michael D. LeClaire, Karina L. Bursch, Brian C. Smith, Madapura M Pradeepa

## Abstract

BRD4 is best known as a transcriptional co-activator, yet heterozygous loss-of-function variants cause craniofacial and neurodevelopmental abnormalities through unclear mechanisms. Using human embryonic stem cells and neural organoids as an in vitro model of human embryonic brain development, we show that BRD4 represses polycomb-regulated developmental and neuronal genes. BRD4 co-occupies bivalent promoters with PRC1.6 and directly interacts with PCGF6 and RING1B through its C-terminal domain. H3K23ac contributes to BRD4 recruitment through BD2, linking acetylated chromatin recognition to polycomb-associated repression. Acute BRD4 depletion alters PRC1.6 occupancy, reduces ubiquitination of H2AK119 and rapidly increases developmental gene transcription. In neural organoids, BRD4 BD2 mutations alter neuronal cell composition and increase expression and chromatin accessibility associated with neuronal transcription-factor programmes. These findings establish BRD4 as a context-dependent chromatin regulator that prevents premature activation of developmental genes and provide mechanistic insight into BRD4-associated neurodevelopmental disorders.

## Introduction

BRD4 is best known as an acetyl-lysine reader that promotes transcription at active promoters and enhancers. However, heterozygous loss-of-function variants in BRD4 cause craniofacial and neurodevelopmental abnormalities, suggesting that its established co-activator functions do not fully explain its role during development. BRD4 belongs to the bromodomain and extra-terminal domain (BET) family, together with BRD2 and BRD3 ^1^. These BET proteins have two conserved N-terminal bromodomains (BD1 and BD2), which recognise acetylated lysine residues. BET proteins have non-redundant functions in regulating chromatin structure, transcription and DNA repair ^2–4^. BRD4 interacts with multiple protein partners to regulate diverse nuclear processes, including transcriptional pause release, elongation, splicing, DNA repair, and maintenance of genomic stability ^5^. BRD4 is enriched at active enhancers and promoters, and very high BRD4 occupancy is characteristic of super-enhancers ^6–9^. However, emerging evidence suggests that BRD4 contributes to transcriptional repression. The alternatively spliced short isoform (BRD4-S) promotes HIV-1 latency by repressing HIV transcription ^10^. The long isoform (BRD4-L), which contains the *C*-terminal domain (CTD), suppresses the viral transcription ^11^. BRD4 also interacts with multiple proteins involved in chromatin compaction and transcriptional repression, including polycomb repressive complex 2 (PRC2) subunit embryonic ectoderm development (EED) ^12^ and the H3K9 methyltransferase G9a ^13^.

Developmental genes are maintained in a silent or poised state by polycomb complexes, allowing their activation at the appropriate stage of differentiation. The *Drosophila* ortholog of BRD4 also interacts with polycomb repressive complex 1 (PRC1) which deposits H2AK119ub1 (hereafter, H2Aub1) ^14^. These studies underscore the complex interplay between BRD4 and repressor complexes in fine-tuning gene expression programmes in a context-dependent manner. PRC2, which deposits H3K27me3, is relatively conserved between Drosophila and vertebrates, whereas vertebrate PRC1 has diversified into multiple canonical and variant complexes with distinct subunit compositions and specialised functions ^15^. Among variant PRC1 complexes, PCGF6-containing PRC1.6 is distinguished by the MGA–MAX and E2F6 DNA-binding modules and represses developmental and germ-cell transcriptional programmes ^16^.

Bivalently marked developmental gene promoters contain both active and repressive chromatin features. Although H3K27ac is generally excluded from these regions, other histone acetylations, including H3K23ac, remain enriched ^17^. Whether these acetylation marks recruit BET proteins to polycomb-regulated chromatin, and how they contribute to gene regulation at these promoters, remain unknown. The central nervous system (CNS) is particularly susceptible to perturbations of histone modifications including acetylations. Mutations in genes involved in histone acetylation pathways lead to diverse neurodevelopmental and neurological disorders ^18–22^. BRD4 plays a key role in neuronal activation and transcriptional responses during memory formation ^23^. We previously demonstrated that mutations in BRD4 cause a neurodevelopmental disorder (NDD) with an atypical Cornelia de Lange Syndrome (CdLS) phenotype ^19^. mESCs harbouring a CdLS mutation in the second bromodomain (BD2) of BRD4 exhibit defective DNA damage signalling and repair ^24^. However, the mechanism by which mutations in BRD4, a general transcriptional coactivator, cause specific phenotypes in neurodevelopmental disorders remains poorly understood.

Here, we aimed to define the roles of BRD4 in regulating neuronal chromatin and gene expression using human embryonic stem cells (hESCs), neurons, and unguided neural organoids (UNOs). Using acute degradation combined with transcriptomics and chromatin profiling, we show that H3K23ac promotes BRD4 recruitment through BD2, while the BRD4 C-terminal domain engages PRC1.6. Disruption of this axis derepresses developmental transcriptional programmes and alters neuronal cell states.

## Results

### BRD4 represses neurodevelopmental genes in stem cells

Genetic knockout, loss-of-function mutations, or prolonged depletion of BRD4 can have pleiotropic consequences. To circumvent the deleterious impacts of long-term BRD4 protein loss, we profiled the effects of short-term BRD4 protein loss in hESCs, hESC-derived neurons, and UNOs using proteolysis-targeting chimaera (PROTAC, ZxH-3-26; hereafter, ZxH) ^25^. Meanwhile, we also generated BRD4-degron-tagged hESCs by knocking in a mutant FKBP and an HA tag at the N-terminus of BRD4 (hereafter BRD4-dTAG) ^26^, which allow for acute degradation of BRD4 upon addition of dTAGv1 (Fig. 1a) ^26^. Together, we investigated the effects of loss-of-function mutations in the BD2 domain of BRD4 in hESCs, mESCs, neurons, and UNOs. These complementary approaches enabled us to distinguish rapid transcriptional responses to BRD4 loss from secondary consequences of prolonged depletion.

**Figure 1:**
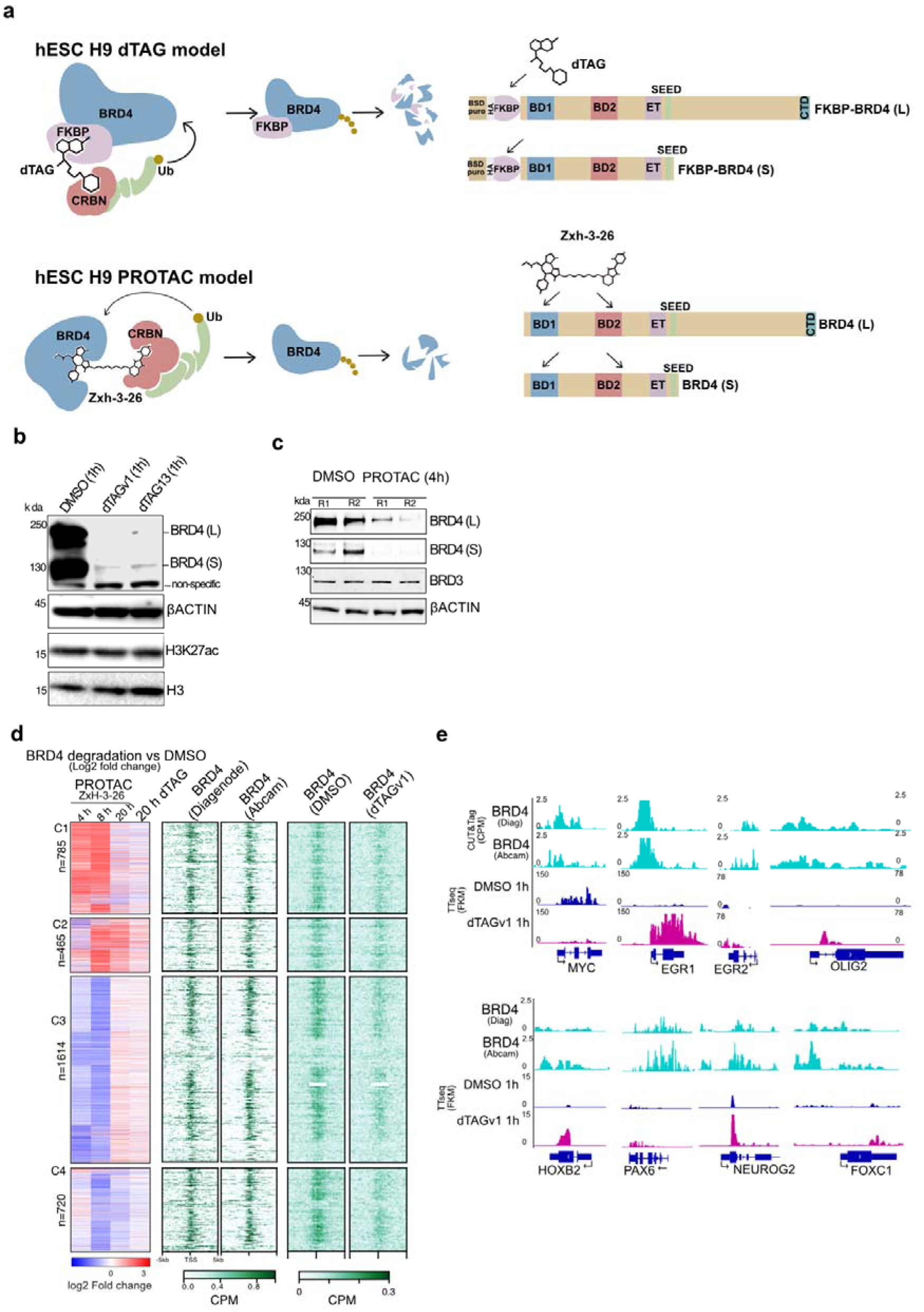
BRD4 degradation upregulates bivalently marked neuronal genes: **a**) Illustration showing the dTAG and PROTAC approaches used for acute BRD4 degradation followed by transcriptomic analysis. **b and c**) Immunoblot showing depletion of BRD4-L and BRD4-S after 4 hours of ZxH-3-26 treatment (ZxH, BRD4-specific PROTAC) (b); dTAGV-1 and dTAG13 mediated BRD4 degradation in BRD4-dTAG hESCs (c), BRD3 and β-ACTIN serve as controls. **d**) Differential RNA-seq from BRD4 PROTAC treatment for (4 hours, 8 hours, 20 hours) in H9 hESCs and 20 hours of dTAGV-1 treatment in BRD4-dTAG hESCs. Heatmap for log2fold change values across four k-means clusters (C1–C4), indicating similar directional changes at least in two of the ZxH treatment time points (left). Heatmaps of CUT&Tag counts per million reads (CPM) signal for short and long isoforms (Diagenode antibody) and long isoform (Abcam antibodies) of BRD4 (gene order is maintained). Heatmap for BRD4 CUT&RUN for DMSO and dTAG mediated BRD4 depletion (1hrs) (right). List of DEGs in Source Data 2. **e**) Genome-browser visualisation of CUT&Tag for BRD4 along with average TTseq signal (n=2 replicates), performed 1 hour after DMSO and dTAGV-1 treatment in BRD4-dTAG hESCs at representative neuro-developmental genes.

Four hours of ZxH treatment in H9 hESCs efficiently depleted both long and short isoforms of BRD4 without affecting BRD2 and BRD3 (Fig. 1b and Extended Data Fig 1). hESCs retained their pluripotency characteristics within four hours of ZxH treatment and 48 hours after drug washout (Extended Data Fig. 1). Similarly, 1-hour dTAGV-1 or dTAG13 treatment in BRD4-dTAG hESCs resulted in near complete degradation of BRD4 (Fig. 1c). We then performed RNA-seq in H9 hESCs treated with ZxH or DMSO for 4, 8 and 20 hours, and in BRD4-dTAG hESCs treated with dTAGV-1 for 20 hours. K-means clustering of genes differentially expressed at two or more time points (adjusted *P* < 0.05) identified four clusters (C1–C4; Fig. 1d). C1 genes were rapidly upregulated after 4 and 8 hours of BRD4 depletion, with a weaker response at 20 hours, whereas C2 genes were predominantly upregulated at 8 and 20 hours. C1 and C2 were enriched for neurodevelopmental, organismal-development and transcriptional processes, and both contained many brain-expressed genes (Extended Data Fig. 2). The reduced magnitude of upregulation at 20 hours may reflect compensatory responses. By contrast, downregulated genes (C3 and C4) were enriched for cell-cycle, pluripotency, DNA-replication and DNA-damage-response pathways (Extended Data Fig. 2).

CUT&Tag using two independent BRD4 antibodies showed enrichment of BRD4 at the TSS of DEGs (Fig. 1d and 1e). Furthermore, the marked reduction in BRD4 CUT&RUN signal following 1 hour of dTAG-mediated degradation confirmed the specificity of the BRD4 antibodies (Figs. 1d and 1e). To assess whether BRD4 directly repress transcription, we performed transient transcriptome sequencing (TT-seq) after 1 hour of dTAG-mediated BRD4 degradation. MYC transcription decreased, consistent with the role of BRD4 in regulating transcription of MYC ^19^. Several neuronal and developmental genes including EGR1, EGR2, OLIG2, HOXB2, PAX6, NEUROG2 and FOXC1 showed BRD4 binding and increased level of nascent transcripts (Fig. 1e). To test reversibility, BRD4 was degraded with dTAGV-1 for 3 hours and subsequently restored by Shield-1 treatment, which prevents further degradation ^27^. BRD4 restoration reversed the upregulated EGR1 and LMO2 and downregulated SOX2 (Extended Data Fig. 3a). Together, these findings support a direct role for BRD4 in repressing a subset of developmental and neuronal genes in hESCs.

### BRD4 co-occupies and regulates PRC1.6 and bivalent promoters

To investigate how BRD4 represses neuronal and developmental genes, we performed CUT&RUN for BRD4-associated factors in H9 hESCs. ChromHMM analysis confirmed BRD4 enrichment at active promoters and enhancers but also revealed occupancy at PRC-repressed and bivalent promoters marked by H3K4me3 and H3K27me3 (Extended Data Fig. 3b) ^28,29^.

Cistrome analysis showed that BRD4 CUT&Tag peaks from hESCs were similar to MAX, and WDR5 (Extended Data Fig. 3c). BRD4 peaks also correlated and overlapped with PRC components and associated histone marks EED, H2Aub1 (Fig. 2a; Extended Data Fig. 3d). Together with H3K14ac and H3K23ac that are known to be enriched at both active and bivalent promoters ^17^. BRD4 showed enrichment not only at active promoters but also at PRC1.6-, PRC2- and bivalent promoters (Fig. 2b). Notably, BRD4 occupancy at PRC1.6 is also detected HEK293 ChIP-seq datasets (Extended Data Figs. 3e and 3f).

**Figure 2:**
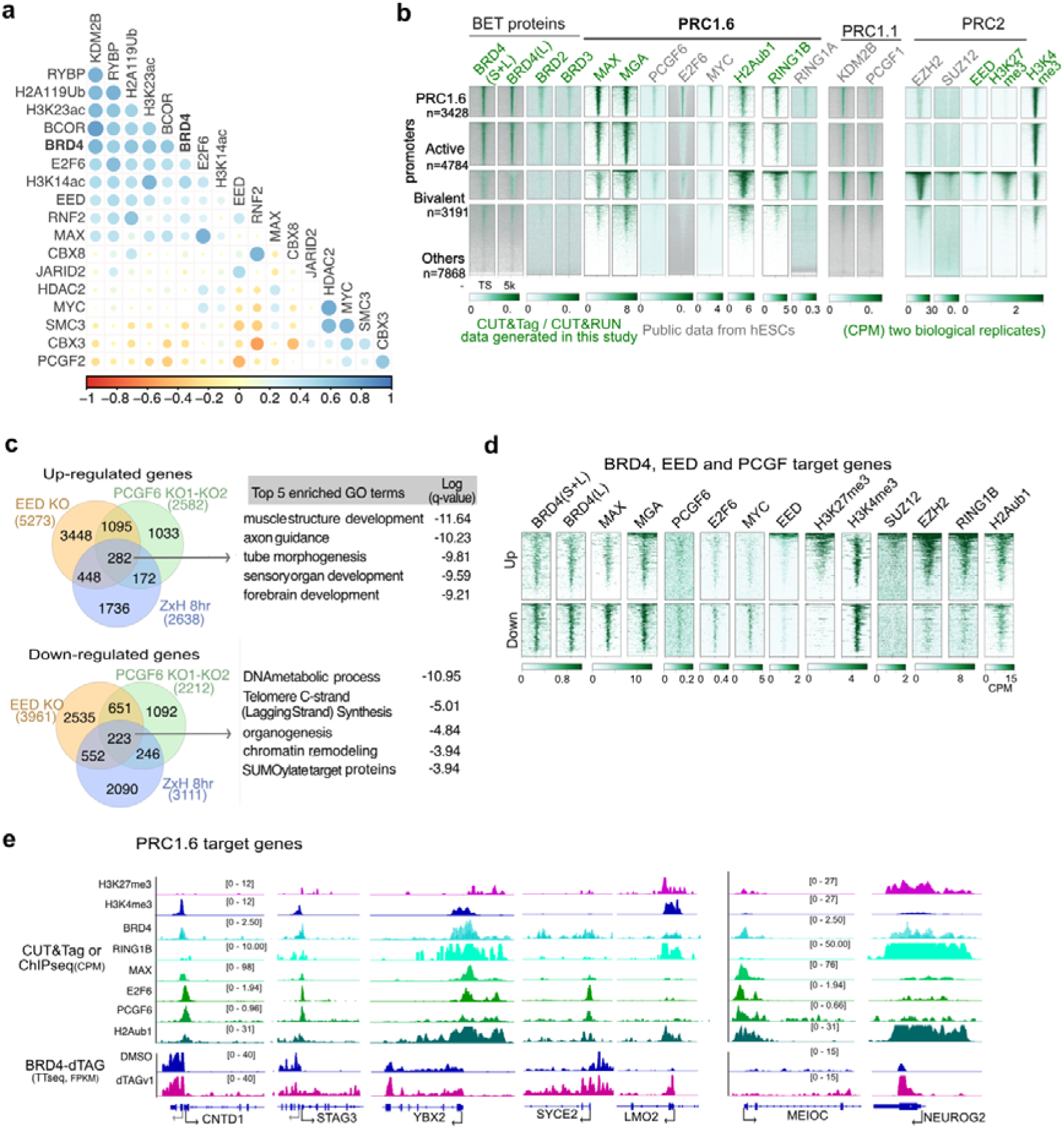
BRD4 occupies and represses PRC target genes. **a)** Pairwise peak intersection correlation plot for chromatin modifications and transcription factors. Values indicate Kendall correlation coefficient between the peak sets ranging from –1 (anti-correlated, red) to 1 (complete correlated, blue) clustered based on principal component. **b**) Heatmaps of CUT&Tag counts per million reads (CPM) for BRD4 (Diagenode antibody), BRD4 (Abcam antibody), H3K27me3, H3K4me3, H3K27ac, CUT&RUN for BRD2, BRD3, EED, MAX, RNF2 and H2Aub1 and; ChIPseq data for PCGF6, MGA, E2F6, MYC, RING1A, KDM2B, PCGF1, EZH2, SUZ12. Clustered based on enrichment of PRC1.6 components, active (H3K4me3), bivalent (H3K27me3+ & H3K4me3+), and other gene promoters. **c**) Venn diagrams and Metascape functional annotations (right) of significant (p-adjusted < 0.05) upregulated (n=282) and significant (p-adjusted < 0.05) downregulated (n=223) genes following 8 hours of ZxH-mediated BRD4 degradation and in two PCGF6 knockout (KO) human pluripotent stem cell lines (data from ^30^) and EED KO ^31^. Details of Metascape analysis in Source Data 2. **d**) Similar to (b), but for commonly up and down regulated genes (identified in panel c) in BRD4 degradation, PCGF and EED KO hESCs. **e**) Like Fig. 1e, but BRD4 signal from two antibodies is merged, also includes histone marks and chromatin proteins. All public datasets used in the study are listed in Extended Data Table 1.

We next aimed to determine whether BRD4 contribute to regulation of PRC1.6 and EED target genes. DEG analysis of publicly available PCGF6- and EED-knockout hPSC RNAseq together with our BRD4 PROTAC showed overlap of 282 up- and 223 down-regulated genes (Fig. 2c) ^30,31^. Upregulated genes are enriched for GO terms associated with nervous system and brain development, while down regulated genes are enriched for DNA metabolic processes (Fig. 2c). Upregulated gene promoters showed high levels of PRC1 and PRC2 proteins and histone modifications deposited by these complexes (H3K27me3 and H2Aub1) compared to downregulated gene promoters (Fig. 2d). These findings identify a shared set of BRD4-, PCGF6- and EED-regulated genes and suggest that BRD4 acts within the polycomb-regulated chromatin environment to restrain transcription of developmental genes.

TTseq data showed increased transcription of known PRC1.6 target genes, including NEUROG2, YBX2, MEIOC, CNTD1 and SYCE2 (Fig. 2e) ^16^. These genes are enriched for BRD4, MAX, RING1B, H3K27me3 and H2Aub, suggesting the direct role of BRD4 and polycomb axis in repressing these genes. We also found that approximately 40% of bivalent genes were upregulated within 4 hours of ZxH treatment, increasing to approximately 50% at 8 hours, with most remaining upregulated at 20 hours (Extended Data Fig. 3g). Together, these findings support a model in which BRD4 cooperates with PRC1.6 to restrain developmental and neuronal gene transcription in hESCs.

### BRD4 interacts with PRC1.6 complex and PRC2 protein EED

To investigate whether BRD4 is specifically associated with PRC complexes, we first analysed the recently published proximal interactomes of PRC1 (CBX7, RING1B/RNF2, PCGF6) and PRC2 (SUZ12 and EZH2) proteins in mouse ES cells ^32^. BRD4 is enriched in the PCGF6 and RING1B interactome, whilst other BET proteins showed low enrichment or were absent (Fig. 3a). Similarly, analysis of our published BRD4 IP-MS in mouse ESCs also showed enrichment for PRC1.6 and PRC2 components (Extended Data Fig. 4a) ^19^. Analysis of another BRD4 IP-MS from K562 cells revealed enrichment of PRC1.6 complex subunits (RING1B/RNF2, MGA, PCGF6, CBX3 and WDR5) (Extended Data Figs. 4b and 4c) ^33,34^. Notably, no other canonical PRC1 (cPRC1) or noncanonical variant PRC1 (PRC1.1-PRC1.5) complex subunits were detected in these BRD4 interactomes.

**Figure 3.**
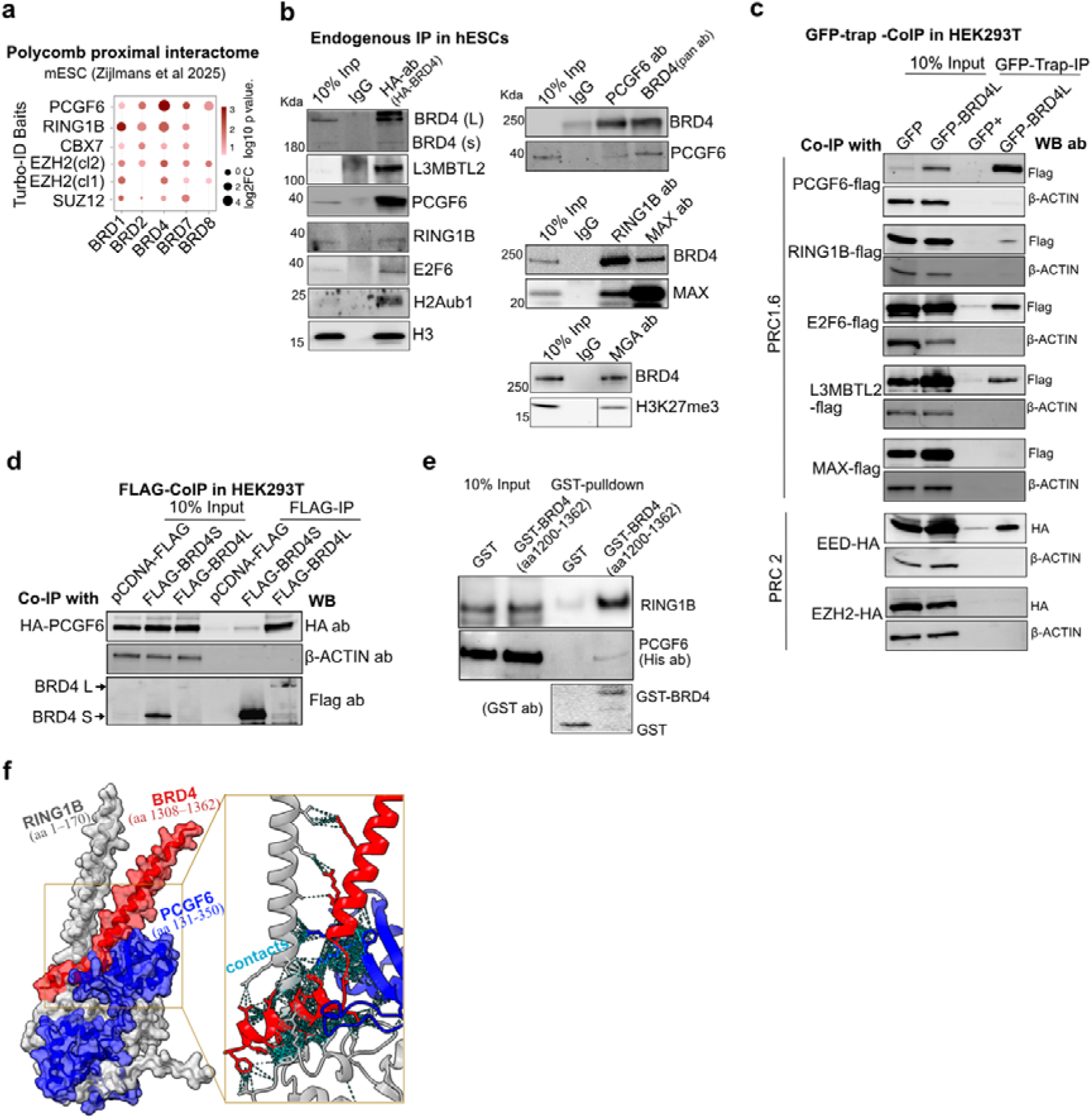
BRD4 interacts with PRC1.6 and EED. **a**) Dot plots showing log2 fold enrichment of BRD proteins in the proximal interactome (Turbo-ID) for PRC1 and PRC2 proteins from mouse embryonic stem cells (mESCs), data from ^32^. The size of the circle represents the log2 fold enrichment in BRD4 IP relative to IgG control. **b)** Immunoblots of HA-IP in HA-FKBP-BRD4 hESCs (left), probed with antibodies against HA (detects BRD4 isoforms), RING1B, PCGF6, E2F6, L3MBTL2, H2Aub1 and H3. Reverse IP using PCGF6, RING1B, MAX and MGA antibodies probed with BRD4, PCGF6, MAX and H3K27me3 antibodies (right). **c**) Immunoblots of GFP-trap co-IP of the GFP-BRD4 long isoform (GFP-BRD4L) with Flag-tagged PCGF6, RING1B, E2F6 and L3MBTL2, MAX, and HA-tagged EED and EZH2. Immunoblots for β-ACTIN served as controls. **d**) Flag pulldown for long BRD4L) and short (BRD4S) form of BRD4 with HA-PCGF6, probed with HA and Flag antibodies. **e**) Immunoblots of recombinant PCGF6-His, and RING1B for invitro GST and GST-BRD4 (C-terminal 1200 to 1362 residues) pulldown. **f)** AlphaFold-predicted structure of a complex comprising the BRD4 C-terminal region (long isoform, aa 1308–1362), the C-terminal region of PCGF6 (aa 131–350), and the N-terminal region of RING1B (aa 1–170). Predicted inter-residue contacts within 5.5 Å between the three proteins are shown in cyan. The model has an ipTM score of 0.50 and a pTM score of 0.49.

**Figure 4.**
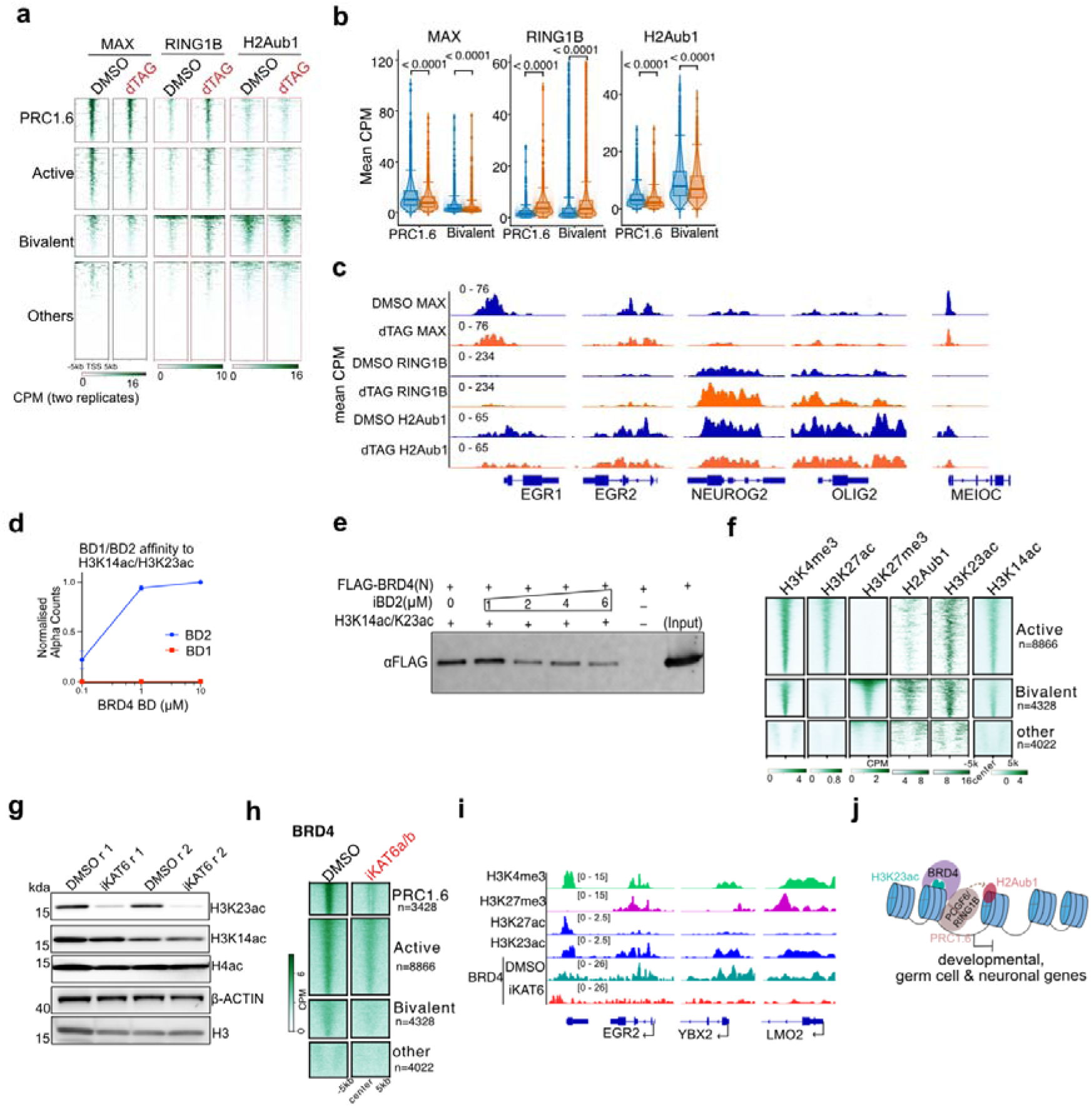
**BRD4 BD2 recognises H3K23ac at bivalent promoters a-c**) Heatmaps (like Fig. 2b, left), violin plots and genome browser tracks for PRC1.6 target genes for MAX, RING1B and H2Aub1 CUT&Tag signal from DMSO and BRD4-dTAG hESCs (20 hours) at PRC1.6 target, active, bivalent and other gene clusters. Centre lines in the violin plot indicate the median, bounds indicate the 25th and 75th percentiles, and whisker limits show 1.5 × interquartile range (p-values from paired t-test with Bonferroni correction). **d)** AlphaScreen counts titration of BRD4-BD1 and -BD2 interaction with H3K14ac/23ac showing that only BRD4-BD2 interacts with H3K14ac/23ac. Normalised average alpha counts of three replicates were set relative to the highest WT. **e)** Immunoblots of H3K14/K23ac peptide pulldown for N-terminal His-FLAG tagged BRD4 (N-terminal 412 amino acids), in the presence of increasing concentration of iBET-BD2 (iBD2). **f**) Heatmap comparing histone marks at active, bivalent and repressed promoters in hESCs as identified in ^37^**. g**) Western blots for H3K23ac, H3K14ac and tetra-acetylated H4 (H4K5/8/12/16ac) 20 hours of DMSO and inhibitor of KAT6a/b (iKAT6a/b) treatment, β-ACTIN and H3 served as controls. **h**) Heatmaps showing BRD4 (CPM) signal at PRC1.6 marked, active and bivalent promoters (like Figs. 2b and 3g) in control (DMSO) and upon iKAT6a/b treatment. **i**) Genome browser tracks for BRD4 CUT&RUN signal in DMSO and iKAT6a/b treated hESCs. **j**) Working model displaying H3K23ac-BRD4-PRC1.6 axis in repressing developmental, germ-cell and neural genes.

Endogenous immunoprecipitation in hESCs confirmed that BRD4 associates with the PRC1.6 components PCGF6, RING1B, E2F6, L3MBTL2, MAX and MGA (Fig. 3b). In HEK293T cells, overexpressed long isoform of BRD4 also interacted with PCGF6, RING1B, E2F6 and L3MBTL2, but not MAX. BRD4 additionally associated with EED, but not EZH2, consistent with previous findings (Fig. 3c) ^12^. Isoform-specific co-IP showed that PCGF6 interacts with long isoform of BRD4 but not short isoform, suggesting the interaction could be mediated through C-terminal domain (CTD) unique to long isoform. Recombinant pulldown assays further supported direct binding of the BRD4 CTD to PCGF6 and RING1B (Fig. 3e), while AlphaFold modelling predicted a potential interface between these regions (Fig. 3f). Notably same domain also binds P-TEFb which facilitate transcriptional elongation^35^.

### BRD4 modulates PRC1.6 occupancy and activity at target sites

CUT&Tag profiling 20 hours after BRD4 degradation showed reduced MAX occupancy and H2Aub1 levels at PRC1.6-regulated promoters, whereas RING1B increased at PRC1.6 and bivalent promoters (Figs. 4a-c). EED occupancy remained largely unchanged (Extended Data Fig. 3h). These findings suggest that BRD4 mediates PRC1.6 occupancy and activity, with increased RING1B but reduced H2Aub1 indicating altered complex assembly or redistribution. Further work will be required to determine how BRD4 regulates occupancy of PRC1 complexes and activity and at active and bivalent promoters.

### BRD4 recognises H3K23ac at active and polycomb regulated promoters

H3K27ac is largely absent from polycomb-repressed and bivalent chromatin because it is antagonised by H3K27me3. However, bivalent nucleosomes can recruit KAT6b, which acetylates H3K23 in mESCs ^17^. Consistent with this, KAT6b was enriched in BRD4 IP–MS, also known to interact with RING1B ^17^. We therefore tested whether H3K23ac recruit BRD4 via its BDs. AlphaScreen assays showed that BRD4-BD2, but not BD1, bound strongly to H3K14ac/K23ac peptides (Fig. 4d; Extended Data Figs. 4d and 4f). Histone peptide pulldown confirmed binding of the BRD4 N-terminus to H3K14ac/K23ac, which was reduced by the BD2-selective inhibitor iBET-BD2 (Fig. 4e) ^36^. iBET-BD2 also reduced BRD4–H3K23ac and BRD4–H3K27me3 PLA foci, while H3K14ac and H3K23ac were enriched in BRD4 IP (Extended Data Figs. 4g and 4h). Like mESCs, bivalent promoters in hESCs were enriched for H3K14ac and H3K23ac, but not H3K27ac in hESCs (Fig. 4f; Extended Data Fig. 4b). Treatment with inhibitor of KAT6a/b (PF-9363) selectively reduced H3K23ac, without detectable changes in H3K14ac or tetra-acetylated H4 (H4K5/8/12/16ac) (Figs. 4g). This was accompanied by reduced BRD4 occupancy at active, bivalent and PRC1.6-regulated promoters, suggesting that H3K23ac contributes to BRD4 recruitment at these sites (Figs. 4h and 4i). Together, these findings support a model in which H3K23ac promotes BRD4 recruitment through BD2, while the BRD4 C-terminal domain engages PRC1.6 at polycomb-regulated promoters (Fig. 4j).

### BRD4 BD2 mutations lead to upregulation of developmental and neuronal genes in mouse and human ESCs

To test how CdLS-associated bromodomain variants affect histone recognition, we compared recombinant BRD4-BD1 N145G and BRD4-BD2 Y390C and Y430C with their wild-type counterparts using AlphaScreen assays (Extended Data Figs. 4d–f). All three variants showed reduced binding to tetra-acetylated H4, indicating that these substitutions impair recognition of at least some acetylated histone substrates. In contrast, BRD4-BD2 Y430C retained binding to H3K14ac/K23ac. Thus, Y430C may not cause a general loss of BD2 function or abolish recognition of acetylation at bivalent promoters. Instead, it appears to alter substrate selectivity. Because acetylated H4 and H3 lysines are enriched in different chromatin contexts ^17,38–40^, this selective defect could redistribute BRD4 across the genome. In vivo, this effect may be further shaped by multivalent interactions with other histone marks, DNA and protein partners.

We then asked whether BRD4 binding to PRC sites is conserved in mouse, and BRD4 mutations leads to deregulation of polycomb repressed genes. We reanalysed our previously published nascent RNA-seq (4su-seq) data from mESCs carrying a CdLS mutation in BD2 of BRD4 (Y430C) ^19,35^, together with public ChIP-seq datasets (Extended Data Table 1). This analysis revealed BRD4 enrichment at PCGF6 peaks but not at peaks of other PRC proteins (Fig. 5a). Furthermore, CdLS mutation in mESCs leads to increased transcription of known PRC1.6 target genes, including meiotic genes (Fig. 5b and 5c). These findings demonstrate conserved role of BRD4 in repressing PRC1.6 target genes in mouse and human ESCs.

**Figure 5:**
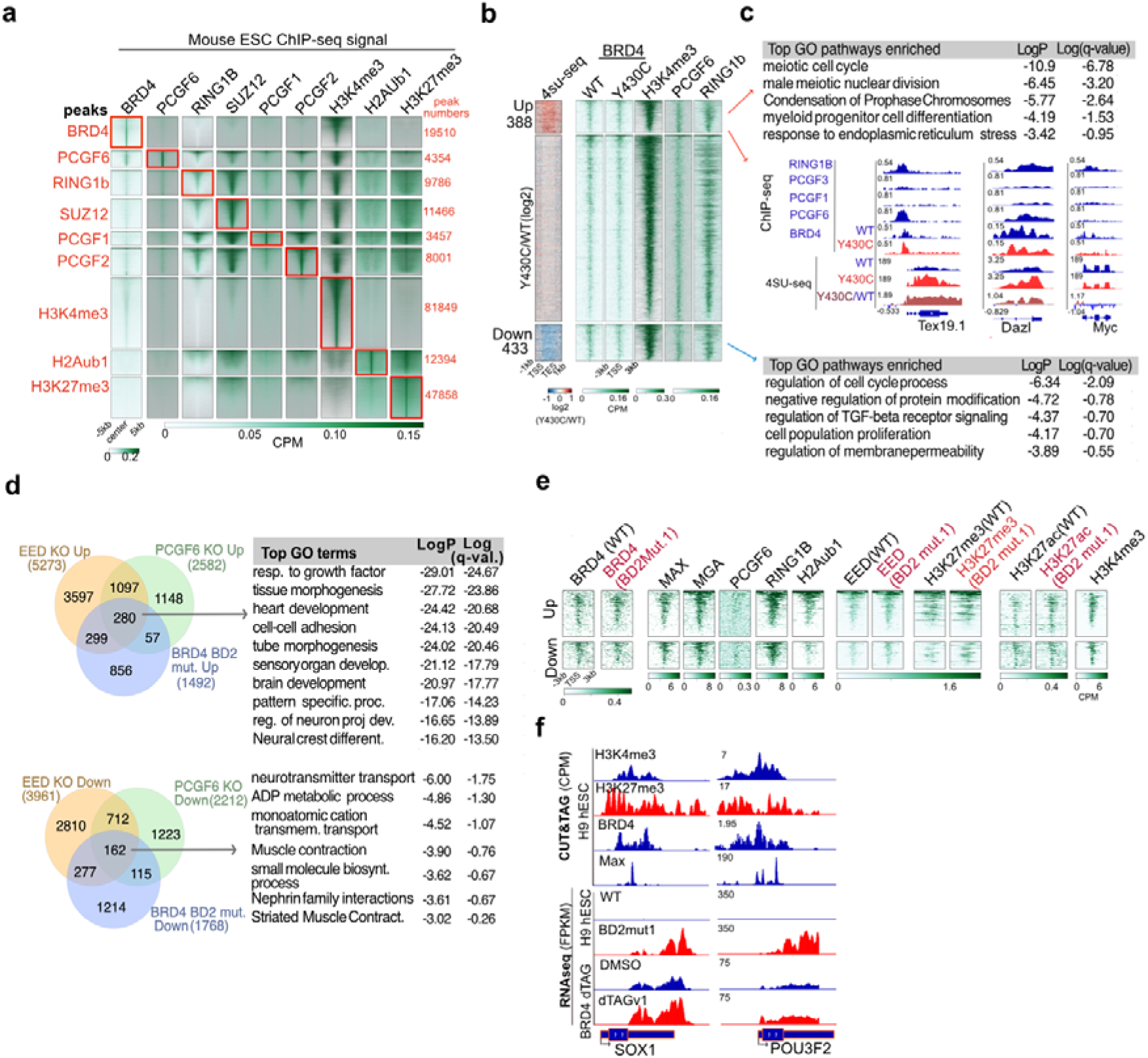
BRD4 BD2 mutations lead to upregulation of PRC1.6 target and neuronal genes. **a**) Heatmap comparing the ChIP-seq signal (CPM) for BRD4 (GSE130659), PCGF6, RING1B, SUZ12, PCGF1, PCGF2, H3K4me3, H2Aub1 and H3K27me3 (GSE122715) respectively at their peaks highlighted in red in mESCs. signal (CPM) for WT and BRD4 BD2 mut1 at protein-coding genes and active enhancers of hESCs. **b**) Heatmap showing 4su-seq log2-ratio (Y430C vs WT, GSE130659) at genes (scaled from TSS-TES) harbouring PCGF6 peaks (from a) clustered using k-means clustering. ChIP-seq signal for BRD4 in WT, Y430C (GSE130659), PCGF6, H3K4me3 and RING1B (GSE122715). c) Top representative GO-terms inferred from Metascape annotations for genes up (red, cluster1) and down (blue, cluster3) as in 4su-seq heatmap in b. Genome browser snapshots showing read pileup of RING1B, PCGF1, PCGF3 (GSE122715), BRD4 (WT, Y430C) and 4su-seq in WT, Y430C and Y430C/WT (GSE130659) at Tex19.1, Dazl and Myc. d) Venn diagrams and Metascape functional annotations (right) of significant (p-adjusted < 0.05) upregulated (top venn diagram in center, n=280) and significant (p-adjusted < 0.05) downregulated (bottom venn diagram in center, n=162) genes in BRD4-BD2 mutant1 and in two PCGF6 knockout (KO) human pluripotent stem cell lines (data from ^30^) and EED KO (PMID 28939884) compared to respective WT or DMSO. e) Heatmap comparing BRD4, EED, H3K27me3, H3K27ac in WT and BRD4 BD2-Mut1 along with MAX, MGA, PCGF6, RNF2, H2Aub1 and H3K4me3 at TSSs (±5kb) of genes common upregulated (top) or downregulated (bottom) identified in panel d. **f**) Genome browser snapshots for CUT&Tag (BRD4, H3K4me3, H3K27me3 and MAX) and RNAseq (WT, BRD4 BD2-Mut1 and DMSO, BRD4-dTAGv1, 3 replicates each).

We aimed to generate CdLS mutation (Y430C) in hESCs (H9) using CRISPR and a repair template, which resulted in homozygous deletion at amino acid 429 (BRD4^C429Δ^, hereafter BD2mut1) due to low homologous recombination rate in hESCs (Extended Data Figs. 4i and 4j). Although these engineered deletions do not reproduce the intended Y430C substitution, they provide independent models of impaired BD2 function. Consistent with the Y430C mutation in mESCs, BRD4-BD2mut1 exhibited reduced occupancy at enhancers and promoters in hESCs (Extended Data Fig. 4k). We found overlap between significantly (p-adjusted <=0.05) upregulated (n=280) and downregulated (n=162) genes in BD2mut1 and those upregulated in PCGF6 and EED KOs, which are enriched for developmental and neuronal genes (Fig. 5d). Chromatin profiling data showed that commonly upregulated and downregulated genes are bound by PRC1.6 components and BRD4; BD2mut1 showed no major effect on the BRD4, EED, H3K27me3 and H3K27ac levels at these promoters.

Commonly upregulated genes showed higher levels of MGA, RING1B, H2Aub1 and H3K27me3 than downregulated genes (Figs. 5e, 5f). Taken together, we conclude that genetic mutations or acute BRD4 depletion led to upregulation of PRC1.6 and EED target genes.

### Acute BRD4 degradation leads to upregulation of neuronal genes in UNOs

We probed the roles of BRD4 in invitro human embryonic brain development model by generating UNOs from H1-hESCs using a well-established protocol ^41,42^. Neurogenesis is known to begin at day 16 in UNOs and to reach highly divergent neuronal lineages around day 42, along with proliferating neural rosettes ^43^ (Figs. 6a and 6b). To identify genes regulated by BRD4 in neuronal cell populations, we performed bulk RNA-seq in UNOs treated with ZxH for 20 hrs to induce acute BRD4 degradation. As in hESCs, acute BRD4 degradation led to upregulation of genes involved in synaptic, axonal, and neuronal projections, including several bivalently marked neuronal immediate-early genes (IEGs) (Figs. 6c-6e). These findings suggest that BRD4 and PRC proteins cooperate to maintain precise temporal control of genes involved in nervous system development and neural plasticity. They also likely play a key role in maintaining IEGs in a transcriptionally poised state, which is critical for rapid activation during the early stages of learning or memory formation ^44,45^.

**Figure 6.**
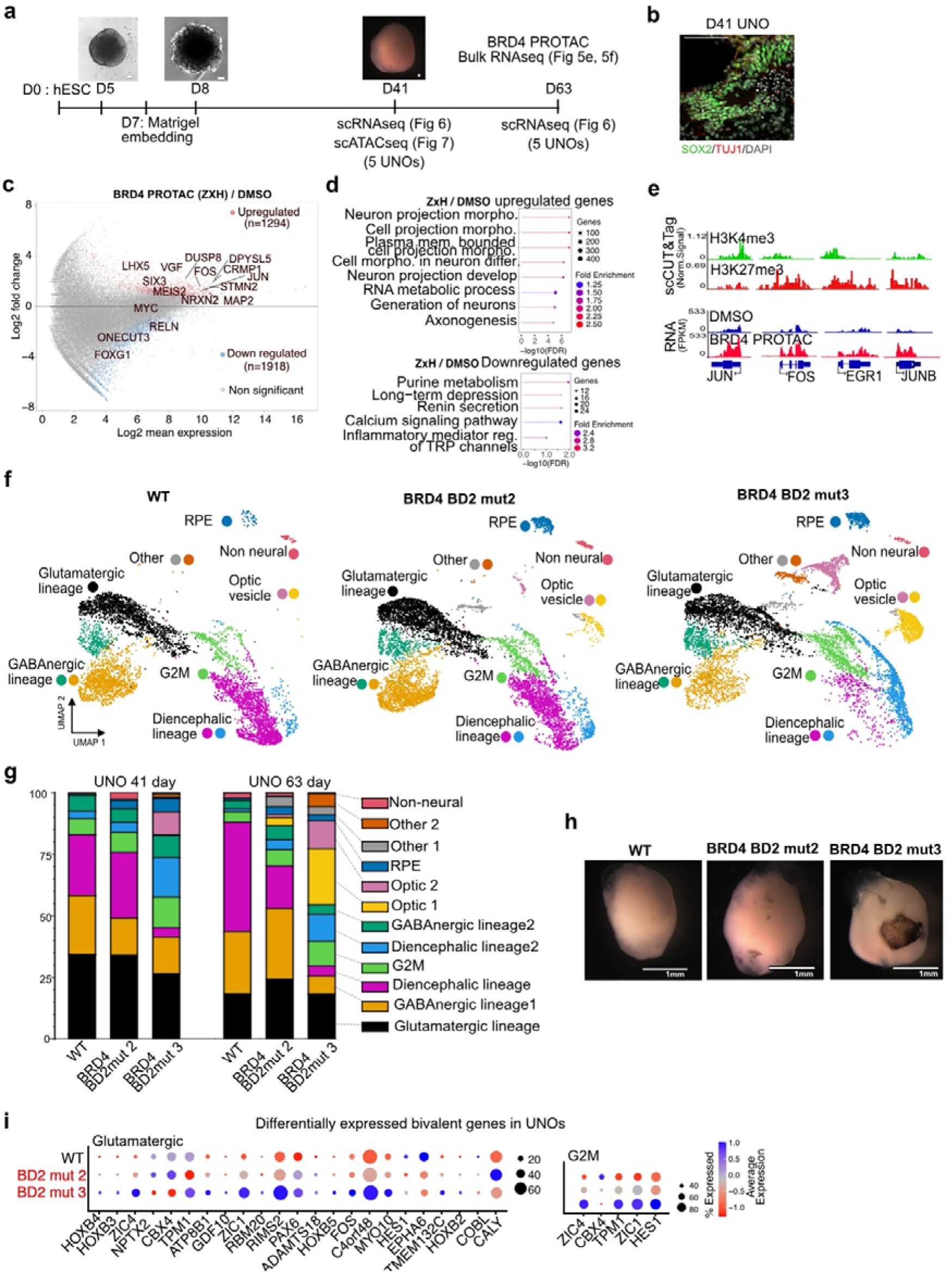
BRD4 BD2 mutations lead to deregulation of bivalent genes and altered cell fate in UNOs. **a)** Schematic representation of the protocol used to generate unguided neuronal organoids (UNOs), with images of UNO WT at 5,8, and 41 days. **b**) Immunofluorescence images of UNOs at day 41 stained for markers of neuronal progenitor (SOX2), post-mitotic early neurons (TUJ1), scale bars: 100 μm. **c**) MA-plot for RNA-seq data illustrating differentially expressed genes in day 41 UNOs following 20 hours of BRD4 PROTAC (ZxH) treatment (n=3 independent organoids). **d)** Gene-ontology (GO) enrichment analyses of up- and down-regulated genes. **e**) Genome-browser tracks for normalised reads at TSS for pseudo bulk scCUT&Tag and bulk RNA-seq for immediate early genes (IEGs) upon 20 h BRD4 PROTAC in UNOs (data from (c)). **f)** UMAP plots stratified by genotype show the annotated cell lineages: WT, BRD4 BD2 mut2 and BRD4 BD2 mut3. Cell clusters are identified by colour, illustrating the contribution of each genotype to specific lineages, such as glutamatergic, GABAnergic, optic vesicle, and RPE. **g)** Stacked bar charts for 41-day and 63-day UNOs, detailing the percentage of cells for each annotated cell type across the WT, BRD4 BD2 mut2 and BRD4 BD2 mut3 UNOs. **h)** Representative bright-field microscopy images of 41-day UNOs, Scale bar=1mm (rest of the images in source file). **i)** Dot-plots showing the average expression level (Z scores) and percentage of cells expressed in glutamatergic, diencephalic-1(pink in UMAP), and diencephalic-2(blue in UMAP), and G2M clusters for bivalent genes that showed significant differential expression in the scRNA-seq data in BRD4-BD2 mut1 and BRD4-BD2 mut2 UNOs.

### BRD4 mutations lead to upregulation of neuronal and bivalent genes and altered cell fate

We next investigated the effects of heterozygous BRD4 loss-of-function mutations (to mimic de novo loss of function variants in CdLS patients) on diverse neuronal cell populations in UNOs as an in vitro embryonic brain development model. For this purpose, we generated H1 hESC clones with two independent heterozygous BD2 mutants. One clone had an in-frame deletion of amino acids 425-430 in one allele (BRD4^425–430Δ/WT^, here on BRD4-BD2mut2) and the second clone had an in-frame deletion of amino acid 429 (BRD4^C429Δ/WT^, here on BRD4-BD2mut3), and an isogenic WT clone (Extended Data Fig. 5a). To investigate the effect of BRD4 mutations on neuronal cell fate and gene expression programme in proliferating neuronal stem cells and in multiple neuronal lineages, we generated scRNA-seq libraries for 41- and 63-day wild-type, BRD4-BD2mut2 and BRD4-BD2mut3 UNOs. To prevent batch effects during library preparation from WT and mutant UNOs (6 UNOs per batch) were multiplexed by antibody-based barcoding at 41 and 63 days. We generated a total of six single-cell libraries totalling 11746 cells at day 41 and 10537 cells at day 63. We further employed the cluster similarity spectrum (CSS) to remove batch effects between time points ^46^.

Dimensionality reduction and embedding of the scRNA-seq data using uniform manifold approximation and projection (UMAP) revealed diverse populations (Fig. 6f and Extended Data Figs. 5b-d) ^47^. Annotation of clusters by comparing the expression of reference markers from each population (Extended Data Fig. 5e) revealed proliferating progenitors (G2M), diencephalic progenitors, GABAergic and glutamatergic lineage, as well as retinal pigment epithelium (RPE), and optic vesicle (Fig. 6f) ^48,49^. Cell type composition analysis showed a reduced diencephalic progenitor cell population 1, and an increased number of diencephalic progenitor cell cluster 2, optic vesicle and RPE cells in the two mutant organoids (Figs. 6f and 6g). This is consistent with the appearance of RPE-like pigmentation in mutant, but not in WT UNOs at day 41 (Fig. 6h). We found enrichment of HOX and ZIC gene families that are known to be regulated by polycomb complexes among genes that are consistently upregulated in both BRD4 mutant UNOs (Extended Data Fig. 6a). It is worth noting that several tested genes upregulated in BRD4 BD2 mutant UNOs are upregulated upon acute BRD4 degradation in hESCs and UNOs.

We then asked whether genes that are upregulated in UNOs with heterozygous BRD4 BD2 mutations are enriched for bivalent modifications. We used publicly available single-cell CUT&Tag (scCUT&Tag) data for H4K4me3, H3K27me3, and H3K27ac from 35-, 60-, and 120-day-old UNOs to generate a list of bivalent chromatin regions in neuronal lineages ^49^. Some but not all these upregulated genes harbour bivalent modifications (Fig. 6i and Extended Data Fig. 6b). Many of these genes are marked with H3K4me3 but lack detectable H3K27me3. The BRD4–PRC1.6 axis likely mediates repression of these genes, as the PRC1.6 complex is known to occupy and repress genes independently of PRC2 ^16^, consistent with the occupancy of BRD4 and PRC1.6 at active genes in hESCs (Fig. 2). We found that several key developmental TFs, which are bivalently marked, are upregulated in neuronal cell lineages (Fig. 6i, and Extended Data Figs. 6a and 6b). These TF families known to play key roles in normal brain development and neuronal cell proliferation and specification. Taken together, our results indicate that BRD4 BD2 mutations leads to deregulation PRC-repressed genes in neuronal cell types.

### Upregulated TF gene families lead to an altered chromatin accessibility landscape in neuronal cells

To define how BRD4 BD2 mutation alters chromatin accessibility, we performed scATAC-seq on day 41 UNOs (6 UNOs per batch), generating 4,236 WT and 4,103 BRD4-BD2mut2 profiles. Consistent with the scRNA-seq data, mutant UNOs showed increased diencephalic and RPE populations (Fig. 7a; Extended Data Figs. 7a,b). Regions gaining accessibility were enriched for C2H2 zinc finger, Homeodomain and bZIP motifs (Fig. 7b; Extended Data Figs. 6a,b and 7c), in line with increased expression of JUN, FOS, EGR1, HOXB2, ZIC1 and ZIC4 (Fig. 6i). Motif enrichment was also cell type specific: diencephalic cells were enriched for bZIP and Homeodomain motifs, glutamatergic cells for T-box and bHLH motifs, GABAergic cells for Homeodomain-POU motifs, and G2M cells for bZIP and SOX motifs (Fig. 7b; Extended Data Fig. 7c). Notably, MGA/MAX recognises T-box and bHLH motifs, supporting reduced PRC1.6-mediated repression following BRD4 perturbation. Altogether these findings show many bivalently marked developmental regulators, including ZIC, PAX, SOX and HOX family genes, were consistently upregulated following acute BRD4 degradation or BD2 mutation, accompanied by increased accessibility of their cognate motifs. Together, these findings indicate that the BRD4-polycomb axis restrains premature activation of developmental transcriptional programmes and helps maintain the balance between cell identity and differentiation.

**Figure 7:**
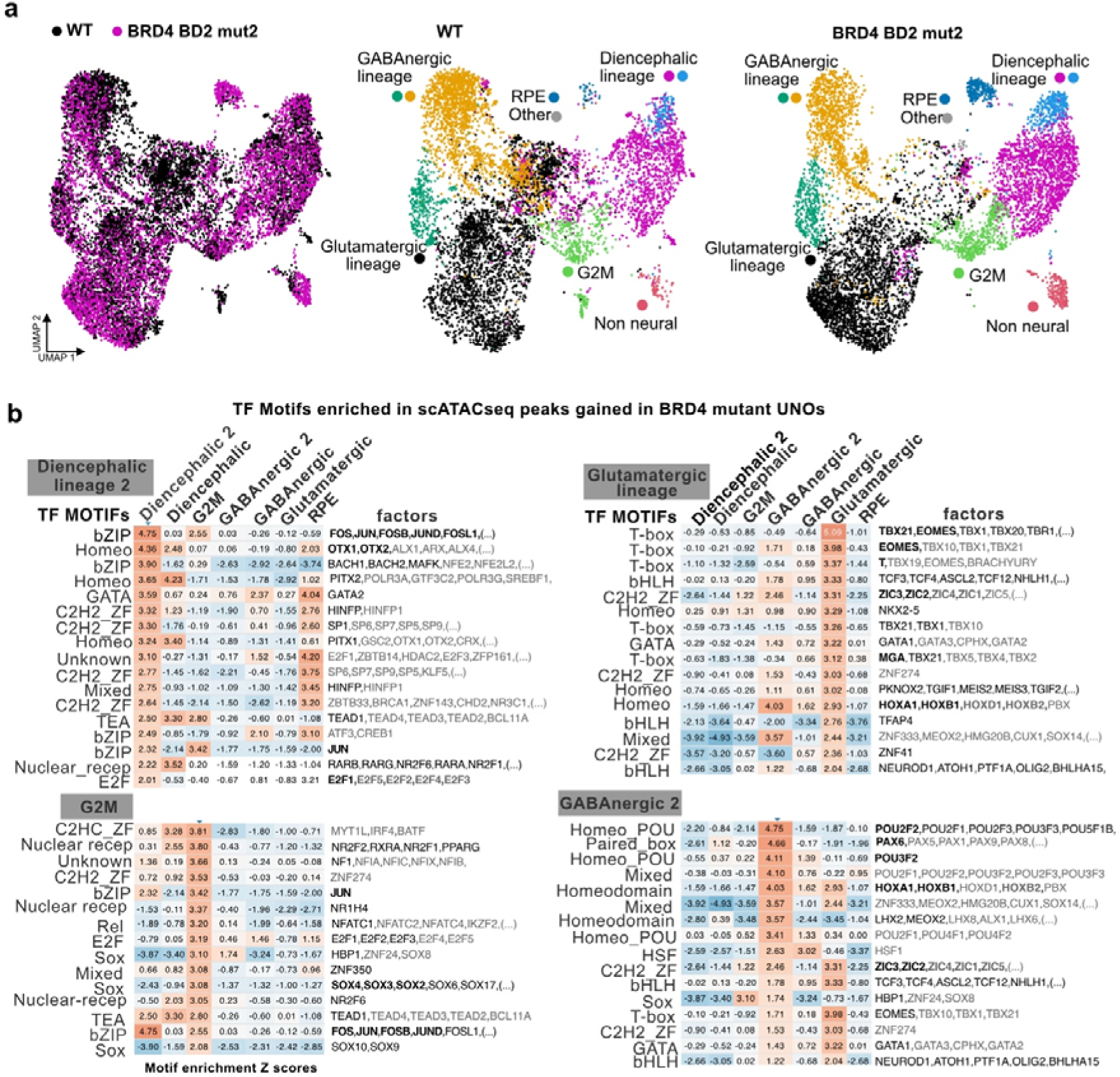
scRNAseq and scATACseq show altered levels of neuronal TF genes BRD4 mutant UNOs. **a)**UMAP plots show the distribution of single-cell ATAC sequencing (scATAC-seq) data clustered by genotypes WT and BRD4 BD2 mut2 and annotated by cell lineage for WT and BRD4 BD2 mut2. **b**) Z-scores (high scores in red and low scores are in blue) showing top transcription factor motifs enriched at diencephalic, glutamatergic, G2M and GABAnergic lineages across scATACseq peaks, which are gained in BRD4 BD2 mut 2 UNO compared to WT control. The complete list of enriched TFs is in the Source Data 3.

## Discussion

Our study identifies an unexpected role for BRD4 in maintaining developmental and neuronal genes in a repressed or poised state. H3K23ac promotes BRD4 recruitment through BD2, whereas the BRD4 C-terminal domain engages PRC1.6, thereby coupling acetylated chromatin recognition to polycomb-associated repression. This resolves the apparent paradox of how a protein best known as a transcriptional co-activator can restrain transcription at selected genomic loci.

### BRD4 interacts with and modulates PRC1.6 activity at developmental genes

Co-occupancy of BRD4 and PRC1.6 components at developmental gene promoters, derepression of PRC1.6-regulated genes following BRD4 depletion, and associated changes in PRC1.6 component occupancy and H2Aub1 levels together provide mechanistic insight into how BRD4 represses developmental and neuronal genes. PRC1.6 can also repress transcription independently of canonical polycomb marks, potentially through associated factors such as L3MBTL2, HP1 and G9A, which may reinforce local chromatin repression ^50–52^. Consistent with impaired PRC1.6 activity, scATAC-seq revealed increased accessibility of bHLH and T-box motifs following BRD4 perturbation. As MGA contains bHLHZip and T-box DNA-binding domains and heterodimerises with MAX, these changes are compatible with reduced MGA–MAX-mediated repression at developmental loci. PRC1.6 also contains L3MBTL2, a chromatin-compaction factor that interacts with BRD4, raising the possibility that disruption of the BRD4–PRC1.6 axis contributes more broadly to increased chromatin accessibility. Direct testing of L3MBTL2, and other PRC1.6 associated proteins such as G9A, HP1 on chromatin compaction and transcriptional repression will be required to establish this mechanism. The combination of increased RING1B occupancy and reduced H2Aub1 argues against simple loss of PRC1 recruitment. Instead, BRD4 depletion may alter PRC1 complex composition, genomic redistribution or catalytic competence. Despite biochemical evidence supporting direct interactions between the BRD4 CTD and PRC1.6 components, the architecture and stoichiometry of the endogenous BRD4–PRC1.6 complex remain to be resolved. Although BRD4 interacts with PRC2 component EED, the functional relevance of the BRD4–EED interaction also remains unclear.

### BRD4 is recruited to polycomb-repressed promoters via binding to H3K23ac

Although PRC-repressed chromatin is depleted of H3K27ac, several other acetylation marks remain enriched, including H3K9ac, H3K14ac, H3K23ac and H3K122ac ^17,28,39,53^. Mechanistically our findings support a model in which H3K23ac promotes BRD4 recruitment via BD2 to polycomb-regulated chromatin, where its CTD engages PRC1.6 components. Because the same BRD4 CTD also binds P-TEFb ^35^, the identity of associated complexes may determine whether BRD4 maintains repression or promotes transcriptional pause release. The CTD interaction with both PRC1.6 and P-TEFb suggests that BRD4 does not possess an intrinsically activating or repressive function. Rather, the local chromatin environment and availability of interacting complexes may determine whether BRD4 promotes pause release or reinforces a poised state.

### BRD4 contributes to repression of developmental and neuronal genes

Acute BRD4 loss rapidly activates developmental transcription factors in hESCs, whereas persistent BD2 perturbation in organoids is accompanied by altered gene-expression programmes, chromatin accessibility and cell-state composition. Patients with BRD4 variants do not display the typical CdLS phenotype, which includes characteristic facial features, growth failure, hypertrichosis of the face and other areas, and radial or limb anomalies ^19^.

However, most patients with BRD4 variants exhibit microcephaly, learning difficulties, intellectual disability, and a global developmental delay phenotype ^19,54^, indicative of nervous system disruptions. BET bromodomain inhibitors and shRNA-mediated depletion of BET proteins in mouse NPCs promotes neurogenesis ^55^. Consistently, we show that loss-of-function mutations in BRD4 or its acute degradation led to the deregulation of key developmental TF families, including HOX, ZIC, FOXG1, SOX, PAX, and SIX. Mutations in these gene families are known to cause various NDD phenotypes ^56–60^. Therefore, many neurodevelopmental and craniofacial abnormalities observed in CdLS cases may stem from the brain-specific TF changes in gene expression and chromatin accessibility resulting from loss-of-function BRD4 variants. The inhibition of BET proteins in a *Xenopus* developmental model also alters pluripotency and neural crest formation ^61^. Similarly, BRD4 depletion also disrupts neural crest differentiation in the mouse model ^4^.

The increased representation of retinal cell states, together with upregulation of retinal regulators such as ONECUT1, SIX3 and PAX6, is consistent with altered specification, expansion or persistence of retinal programmes. These changes are also consistent with ocular phenotypes observed in BRD4-mutant mice and individuals with BRD4-associated disorders ^54,62,63^. However, lineage-resolved experiments will be required to determine whether these changes reflect premature differentiation, altered progenitor allocation or differential cell survival. The proposed pathway is also supported by genetic convergence across neurodevelopmental disorders. Interestingly, mutations in KAT6a and KAT6b, which encode acetyltransferases that deposit H3K23ac, cause NDDs ^22^. Similarly, mutations in regulators that maintain H2Aub1 homeostasis are associated with neurodevelopmental and autism-spectrum disorders ^64,65^. Altogether, our findings suggest a unifying mechanism whereby disruption of the H3K23ac–BRD4–H2Aub1 axis leads to inappropriate regulation of developmental gene expression.

## Limitations

Although the BD2 deletions perturb BRD4 function, they do not precisely reproduce the patient-associated Y430C variant. As Y430C retains H3K14ac/H3K23ac binding in vitro but shows reduced binding to tetra-acetylated H4 in vitro, BD2 mutations may alter BRD4 distribution across acetylated chromatin rather than abolish recruitment to bivalent promoters. In addition, altered cell-state proportions in neural organoids support a developmental effect but do not establish in vivo lineage trajectories.

In summary, BRD4 acts as a context-dependent regulator of developmental genes by coupling recognition of H3K23ac through interactions with PRC1.6. Future work should determine how BRD4-associated repressor and transcriptional elongation complexes are exchanged during gene activation and whether this mechanism operates across additional developmental lineages. These findings provide a mechanistic framework linking histone acetylation, polycomb regulation and neurodevelopmental disease.

## Online Methods

### hESC culture

H9 (WiCell) and H1 hESCs (WiCell) were cultured in mTeSR+ (Stemcell Technologies Cat.# 100-0276) on Matrigel hESC-Qualified Matrix (Corning Cat.# 354277). Cells were split at a ratio of 1/10-1/20 with ReLeSR (Stemcell Technologies Cat.# 100-0484) or Accutase (ThermoFisher Scientific Cat.# 00-4555-56) if the experiment required single-cell splitting. Cells were treated with 100 nM of BRD4 PROTAC (Cambridge Bioscience Cat. # T17297 ZxH-3-26) for 4 to 20h, 5 µM of KAT6a/b inhibitor (Cambridge Bioscience Cat.# HY-132283 PF-9363) for 20h (tested using western blots Tetra-acetylated H4 (Millipore Cat# 06-866, Lot 2282163), H3K14ac (Abcam, Cat# AB203952) and 1 µM of iBET-BD2 (Cayman Chemical, Cat.# Cay31766-1) for 16h. Cell cultures were routinely tested for mycoplasma contamination by semi-quantitative PCR.

### Generation of BRD4 mutant hESCs

We generated our BRD4 mutant hESCs from H9 and H1hESC. We designed gRNA targeting aa430 in the BD2 domain. We use the Nucleofector (Lonza, Nucleofector 4D) with programme number CB150. 0.2 million cells were nucleofected with an RNP complex containing Cas9 (IDT Cat# 1081060) and gRNA (AGGGTTGTACTTATAGCAGT), both from IDT, and P3 Primary Cell 4D Nucleofector X Kit (Lonza, Cat# V4XP-3032). Colonies were manually picked and then sequenced for genotyping.

### Generation of BRD4 dTAG hESC line

We generated H9 hESC with a degron tag (FKBP) at the N-terminus of BRD4 by transfecting RNP containing Cas9 and gRNA targeting the N-terminus of BRD4, along with plasmids containing microhomology for the N-terminus of BRD4. H9 hESCs were nucleofected to H9 hESCs using the Nucleofector 4D (Lonza, Nucleofector 4D with programme CB-150) device and P3 Primary Cell 4D Nucleofector X Kit (Lonza, Cat# V4XP-3032) with RNP containing CAS9 (IDT) and guideRNAs ATGTCTGCGGAGAGCGGCCC (targeting N-terminus of BRD4, IDT), and GCATCGTACGCGTACGTGTT (targeting PITCH, IDT) along with two BRD4 N-terminal dTAG constructs, one expressing puromycin resistance (pCRIS-PITChv2-dTAG-BRD4-puroR, Addgene Cat# 91793) and blasticidin resistance (pCRIS-PITChv2-dTAG-BRD4-blastR, Addgene Cat# 91792). Plasmids were a gift from James Bradner and Behnam Nabet ^26^. Targeted clones were selected with 0.5 mg/ml puromycin and 2.5 μg/ml blasticidin for 10 days, starting five days after nucleofection. Surviving colonies were genotyped by initial PCR and Sanger sequencing, and the edits were further confirmed by western blotting with an HA antibody (Thermo Fisher Scientific, Cat. #26183) to detect correctly edited BRD4; degradation efficiency was verified by treating them with 500 nM dTAGV-1 and dTAG13. HA-FKBP-BRD4 degradation was rescued by replacing the dTAGv1 containing media with 1uM Shield-1 containing media (Clinisciences Ltd. Cat.# 914805-33-7).

### NSC induction of hESCs and neuronal differentiation

Neural stem cells (NSCs) were derived from WT and BRD4 BD2 mutant H9 hESCs using the manufacturer’s protocol (Life Technologies, MAN0008031) and expanded in neural expansion medium (NEM), which consisted of Neurobasal medium(Thermo Fisher; Cat.# 21103049) and advanced DMEM/F12 (Thermo Fisher; Cat.# 12634010). The media was supplemented with a neural induction supplement (Thermo Fisher; A1647801) and penicillin/streptomycin. NSCs were validated by immunolabelling and used for neuronal differentiation between passages 6 and 10. NSCs were differentiated into forebrain neurons using STEMdiff™ Forebrain Neuron Maturation Kit (Stem Cell Technologies, Cat. # 08605) for 21 days to produce a mixed population of excitatory and inhibitory forebrain-type neurons.

### Unguided neural organoids (UNOs)

Unguided neural organoids were generated using the cerebral organoid kit (Stem Cell Technologies, Cat #08570). Briefly, 9000 cells were plated in a low attachment 96-well plate in EB seeding media, centrifuged for 3min at 400 × g, and then topped up with EB formation media on days 2 and 4. On day 5, they are transferred to induction media for 2 days. At day 7, embryoid bodies were embedded in 15 μL of Matrigel basement membrane matrix (Corning, Cat. #354230) and transferred to a low attachment 6-well plate in expansion media. After 3 days in expansion media, they were transferred to maturation media on an orbital shaker at 65 rpm.

For scRNA-seq and scATAC-seq, 5UNO samples were pooled and processed using a papain-based dissociation kit (MACS, Miltenyi Biotech, Cat. # 130-092-628). Briefly, UNOs were cut in half with a scalpel blade and processed with 2 mL of preheated papain solution for 15 minutes at 37 °C. The enzyme mix was added to the UNO pieces and was triturated with wide bore p1000 and regular p1000 pipette tips. They were incubated twice for 10 minutes at 37 °C, with trituration steps in between, using p200 and p1000 pipette tips. Cells were filtered through 30 μm filters, centrifuged, and processed for scRNA-seq and scATAC-seq library preparation using the 10x Genomics Chromium Next GEM Single Cell 3 Kit (Cat.# PN-1000121) and the Chromium scATAC-seq Next GEM Single Cell ATAC v2 Kit (Cat.# PN-1000390). UNOs at days 41 and 63 were processed separately: cells were dissociated from WT and two mutant UNOs, which were labelled with antibody-based hashtag oligos (HTOs), and pooled before loading onto a 10x Genomics chip.

### Immunofluorescence on UNOs

UNOs were fixed in 4% paraformaldehyde at 4°C overnight, washed 3 times with PBS, embedded in OCT after a sucrose gradient, and frozen at –80 °C. Sections were cut with a cryostat LEICA CM1850 at 20μm and placed on microscope slides (Epredia, Cat# J1800AMNZ). Sections were washed with PBS, permeabilised using 0.15%Triton X-100, blocked using 0.15%Triton X-100/2%BSA for 1h, and incubated at 4 °C overnight with primary antibodies anti-SOX2 (Santa Cruz Biotechnology, Cat# sc-365964, 1/1000) and anti-TUJ1 (BioLegend, Cat# 802001, 1/1000). After washing with PBS-0.15 % Triton X-100 three times, the sections were incubated with secondary antibodies conjugated to different fluorophores and DAPI for 2h at RT, then washed with PBS-0.15 % Triton X-100 and mounted with Dako mounting medium. Sections were imaged with a Zeiss LSM 800 confocal microscope.

### Proximity Ligation Assay (PLA)

WT and BRD4 mutant hESCs were seeded in glass-bottom 96-well plates. PLA was performed using the Duolink InSitu Red Starter kit (Mouse/Rabbit) according to the manufacturer’s protocol (Merck, DUO92101) and permeabilized using 0.1% Triton-X-100 for 5min. Antibodies anti-BRD4 (Abcam, Cat.# ab244221, 1/200), anti-H3K27ac (Abcam, Cat. # ab4729, 1/800), anti-H3K27me3 (Millipore Cat. # 07-449, 1/500) and anti-H3K23ac (Merck Cat.# 07-355, 1/500) were used. Plates were imaged in the laser-based InCell high-content imaging system (IN Cell Analyser 6000, GE Healthcare). The images were analysed using INCarta. The data used for comparison are listed in Extended Data 1.

### Bulk RNA-seq in WT and BD2 mutant hESC, and neurons

For WT and BD2 mutant H9 hESCs, NPCs and neurons, total RNA was isolated using NEB Monarch kits; ribosomal RNAs were depleted using the RiboCop kit (Lexogen, Cat.# 144), RNA-seq libraries were generated using the CORALL total RNA-seq V2 kit (Lexogen Cat.# 183) followed by sequencing as 150bp paired-end reads.

### Bulk RNA-seq upon dTAG and PROTAC treatment

RNA-seq libraries upon BRD4 PROTAC (TargetMol Chemicals, Cat. # T17297, ZxH-3-26) in H9 hESCs, and dTAGV-1 treatment in H9 hESC-BRD4-dTAG lines were generated using NEBNext^®^ Poly(A) mRNA Magnetic Isolation Module (NEB, Cat.# E7490L) followed by NEBNext^®^ Ultra^™^ II RNA (NEB, Cat.# E7770L).

### Endogenous Immunoprecipitation

Cells were lysed in native IP buffer (20 mM Tris, pH 8.0, 50 mM NaCl, 0.5% (vol/vol) IGEPAL CA-630, 10% (vol/vol) glycerol) containing protease and phosphatase inhibitors, and incubated on ice for 4 min. Nuclei are centrifuged at 15500 × *g* for 10 min, then resuspended in native IP buffer + Benzonase (Millipore Sigma, Cat.# 707463 final concentration, 1.25 U/μl) for 30 min on ice. Extracts were cleared by centrifugation at 15500 *g* for 10 min at 4°C. The supernatant was precleared with Dynabeads A bead on ice for 5min. The precleared extract was then rotated with 5ug of HA-tag, 1 μg of antibodies anti-BRD4 (Cell Signalling, Cat. # 83375), anti-RING1B (Active Motif, Cat. # 39663), anti-MGA (Antibodies Online, Cat.# ABIN2444597), PCGF6 (Abcepta, Cat# AP18850c-ev), anti-MAX (Proteintech Cat#10426-1-AP) or IGG (ThermoFisher Scientific, Cat. # 500-P00-500UG) for 2 hours at 4 °C before adding the Dynabeads A bead for 30min at 4 °C. After two washes in native IP buffer and three more in IP washing buffer (20 mM Tris, pH 8.0, 125 mM NaCl, 0.5% (vol/vol) IGEPAL CA-630 and protease inhibitors), bound proteins were eluted by heating the beads with 1× Bolt sodium dodecyl sulphate (SDS) loading buffer, and separated on a Bolt™ Bis-Tris Plus, transferred to a nitrocellulose membrane, and immunoblotted with primary antibodies anti-BRD4 (Santa Cruz Biotechnology, Cat. # sc-518158), anti-BRD3 (Active Motif, Cat. # 61489), anti-RING1B (Active Motif, Cat. # 39663), anti-CBX7 (Abcam, Cat.# ab21873), PCGF6 (Abcepta, Cat# AP18850c-ev) anti-MAX (Proteintech Cat#10426-1-AP), anti-H3K27me3 (Millipore, Cat.# 07-449), anti-H3K23ac (Millipore, Cat.# 07-355), anti-E2F6 (Insight Biotechnology Cat.# SAB-33362-1), L3MBTL2 (Proteintech, Cat.# 24435-1-AP), H2Aub1 (Merck Cat# 05-678) and H3K14ac (Active Motif, Cat.# 65701) overnight at 4°C or 1 hour at room temperature on a nutator. After three washes with PBST, membranes were incubated with fluorescence secondary antibodies (StarBright™ Blue 520 Goat Anti-Mouse IgG BioRad Cat. # 12005867, StarBright Blue 700 Goat Anti-Rabbit IgG, BioRad, Cat. # 12004162 and DyLight 800 Anti-Mouse IgM ThermoFisher Scientific Cat. # SA510156) or HRP secondaries antibodies (TidyBlot BioRad Cat. # STAR209PT and Anti-Mouse IgM Invitrogen Cat. # 31440) and Anti-ACTIN hFAB™ Rhodamine Antibody BioRad, Cat. # 12004164). After three washes with PBST, the membranes were imaged using a BioRad imager.

### Co-immunoprecipitation (Co-IP)

Co-IP was performed using the GFP-trap kit according to the manufacturer’s protocol (ChromoTek GFP-Trap® Agarose kit, Cat # gtak), with modifications described below. ∼1.2 million HEK293T cells were reverse transfected with 1 μg of long isoform of GFP-BRD4 (Addgene 65378)^66^, pEGFP (Clontech) plasmids, along with 1μg of pCMV-HA-hEZH2 (Addgene # 24230), pCMV-HA-EED (Addgene # 24231)^67^, pcDNA3-FLAG-E2F6 (Addgene 236441)^68^, pCMV-FLAG-hL3MBTL2 (Addgene # 28232)^69^, pCAG-3xFlag Ring1B-IRES-Puro, pCAG-3xFlag, Pcgf6-IRES-Bsd, pCAG-3xFlag.Max-IRES-Puro ^70^ in 6 well plates using Lipofectamine 3000 reagent (ThermoFisher, Cat# L3000015). 24 hours after transfection, cells were lysed using 200 μl of RIPA lysis buffer (10 mM Tris/Cl pH 7.5, 150 mM NaCl, 0.5 mM EDTA, 0.1 % SDS, 1 % Triton™ X-100, 1 % deoxycholate, 0.09 % sodium azide, supplemented with Benzonase, proteinase inhibitor cocktail and PMSF) for 1 hour, lysate was cleared by centrifugation at 17000xg, supernatant was mixed with 300 μl of dilution buffer (10 mM Tris/Cl pH 7.5, 150 mM NaCl, 0.5 mM EDTA, 0.018 % sodium azide, proteinase inhibitor cocktail and PMSF). After saving 10% of the lysate for input, 450 μl of lysate is incubated with 15 μl of GFP-trap Agarose beads, equilibrated in dilution buffer, in an end-to-end rotor at 4 °C for 1 hour. Beads were washed with wash buffer (10 mM Tris/Cl pH 7.5, 150 mM NaCl, 0.05 % Nonidet™ P40 Substitute, 0.5 mM EDTA, 0.018 % sodium azide) and eluted by heating samples at 95C for 7 min in 1× Bolt sodium dodecyl sulphate (SDS) loading buffer, and separated on a Bolt™ Bis-Tris Plus, transferred to a nitrocellulose membrane, and immunoblotted with anti-Flag (Sigma Aldrich Cat# F7425) or anti HA (Proteintech Cat# 51064-2-AP) antibodies for overnight at 4C. After three washes with PBST, membranes were incubated with fluorescence secondary antibodies (StarBright Blue 700 Goat Anti-Rabbit IgG, BioRad, Cat. # 12004162 and Anti-ACTIN hFAB™ Rhodamine Antibody BioRad, Cat. # 12004164). After three 5-minute washes with PBST, the membranes were imaged using a BioRad imager.

Flag pull-down was performed by co-transfecting Flag-BRD4 short pCMV-FlagBRD4S and long pCMV-FlagBRD4L isoforms ^71^ or pCDNA with Pcgf6-IRES-Puro and pCAG-3xHA ^70^, followed by Flag-M2 magnetic beads under the same conditions as GFP-Trap Co-IP.

### AlphaFold modelling

Structural modelling of the BRD4-PCGF6-RING1B heterotrimeric complex was performed using the AlphaFold 3 server. To optimise structural alignment efficiency and isolate the core interaction interfaces, truncated sequences representing specific domains were used as inputs for modelling: BRD4: The C-terminal 54 amino acids of the long isoform (residues 1309–1362). PCGF6: The C-terminal 220 residues (residues 131–350). RING1B: The N-terminal 170 residues (residues 1–170). The ternary complex was modelled by specifying exactly one copy of each truncated protein chain in the structural pipeline. pLDDT (predicted Local Distance Difference Test): Monitored local, per-residue confidence in the backbone structure. The top-ranked structural model, determined by the highest combined ipTM (0.5) and pTM (0.49) metrics, was selected for further visualisation using UCSF ChimeraX.

### Protein purification and in vitro pulldown

C-terminal BRD4 (BRD4 C-ter, residues 1200-1362) was PCR-amplified and cloned into pGEX4T1. GST-BRD4-Cter, GST-RING1B and GST proteins were expressed in BL21 and purified using Glutathione agarose beads. GST-RING1B (kind gift from Justyna McIntyre, Polish Academy of Sciences, Warsaw) was eluted by thrombin cleavage, resulting in untagged RING1B. PCGF-His was purified using a Cobalt column and eluted with 200mM imidazole. 10 µl of glutathione agarose beads were incubated with purified proteins in an in vitro pulldown buffer (50mM HEPES-KOH pH 7.5, 200mM NaCl, 1% Triton X-100, 10% glycerol, 1mM EDTA, 1mM DTT and Complete Protease inhibitor 1X) for 1 hour at 4 °C. Beads were washed for 3 times in in vitro pulldown buffer and eluted by heating samples at 95 °C for 7 min in 1× Bolt sodium dodecyl sulphate (SDS) loading buffer, and separated on a Bolt™ Bis-Tris Plus, transferred to a nitrocellulose membrane, and immunoblotted with anti-His tag (Sigma Aldrich Cat# H1029) or anti RING1B antibodies (Active Motif Cat. # 39663) for overnight at 4 °C.

### Histone peptide pulldown

1 μg of biotinylated histone H3K14ac/H3K23ac peptide (Cayman Chemicals, Cat. 27520-250ug-CAY) was incubated with 10 μL of streptavidin magnetic beads (Invitrogen 656-01) in 300 μL of binding buffer (50 mM Tris, pH 7.5, 200 mM NaCl and 0.1% NP-40, proteinase inhibitor cocktail) and rotated at room temperature for 30 min. At the same time, FLAG-His tagged BRD4 *N*-terminal domain containing BD1 and BD2 (E49-E460) (MedChemExpress Cat# HY-P7846), iBET-BD2 (Cayman Chemical Cat# CAY31766), or DMSO were added to the binding buffer on ice. Biotinylated H3K14ac/K23ac peptide-bound streptavidin beads were added to the BRD4 in binding buffer and incubated overnight at 4 °C with rotation. The beads were washed with 1 mL of cold binding buffer 3 times with 3 min rotation each. Beads were resuspended in 30 μL of 1x Bolt sample loading buffer and heated at 95 °C for 5 min, then separated proteins using Bolt™ Bis-Tris Plus gels. Followed by western blotting using anti-FLAG antibody to detect BRD4.

### CUT&Run

CUT&Run was performed according to 74 with modifications from the EpiCypher® CUTANA™ CUT&Run Protocol, pAG-Mnase from Epicypher (Cat. # SKU: 15-1116), and Cell Signalling Technology (Cat. # 403665). CUT&Run was performed using 500,000 cells. CUT&Run libraries are prepared using the NEBNext^®^ Ultra™ II DNA Library Prep Kit for Illumina (Cat # E7645L) by following the protocol version 2 of the protocol.io approach adapted for CUT&Run library sequencing (dx.doi.org/10.17504/protocols.io.bagaibse). Primary antibodies used were anti-RAD21 (Abcam, Cat.#ab217678), anti-EED (Proteintech, Cat.# 16818-1-AP), anti-NIPBL (Bethyl, Cat.# A301-778A), anti-Phospho-Rpb1 CTD (Ser5) (Cell Signaling Technology, Cat.# 13523), anti-H3K27me3 (Diagenode, Cat.# C15410195), anti-BRD2 (Abcam, Cat.# ab139690), anti-BRD3 (Active Motif, Cat.# 61489) and anti-H3K4me3 (Millipore, Cat.# 07-473).

### CUT&Tag

CUT&Tag was performed according to the protocol ^73^, with modifications to tissue processing as described below. Experiments were performed in biological duplicates from each cell type. ∼50,000 cells were pelleted by centrifugation for 3 min at 600 × g at room temperature and resuspended in 500μl of ice-cold NE1 buffer (20 mM HEPES-KOH pH 7.9, 10 mM KCl, 0.5 mM spermidine, 1% Triton X-100, and 20 % glycerol and cOmplete EDTA-free protease inhibitor tablet) and were let to sit for 10 min on ice. Nuclei were pelleted by centrifugation for 4 min at 1300 × g at 4°C, resuspended in 500 μl of wash buffer, and held on ice until the beads were ready. The required amount of BioMag Plus Concanavalin-A-conjugated magnetic beads (Polysciences, Cat.# 86057) was transferred into the binding buffer (20 mM HEPES-KOH pH 7.9, 10 mM KCl, 1mM CaCl2 and 1mM MnCl2), washed once in the same buffer, each time placing them on a magnetic rack to allow the beads to separate from the buffer and resuspended in binding buffer. 10 μl of beads was added to each tube containing cells, and the tubes were rotated on an end-to-end rotator for 10 minutes. After a pulse spin to remove liquid from the cap, the tubes were placed on a magnet stand to clear the liquid. The liquid was then withdrawn, and 800 μl of antibody buffer containing 1 μg of primary antibodies was added and incubated at 4 °C overnight in a nutator. Primary antibodies used were anti-BRD4 (Diagenode, Cat.# C15410337), anti-BRD4 (Abcam, Cat.# ab128874), anti-H3K27ac (Abcam, Cat.# ab4729), and H3K23ac (Millipore, Cat.# 07-355), H2Aub1 (Cell signalling Technologies Cat. 8240), H3K27me3 (Diagenode, Cat. # C15200181-10, Lot 001-12), MGA (Antibodies online, Cat.# ABIN2444597), RNF2/RING1B (Sino Biologicals Cat. 202325-T38-100), MAX, (Proteintech Cat# 10426-1-AP). Secondary antibodies (guinea pig α-rabbit antibody, Antibodies online, Cat.# ABIN101961) were diluted 1:100 in dig-wash buffer (5% digitonin in wash buffer), and 100 μL was added to each sample while gently vortexing to dislodge the beads from the sides, then incubated for 60 min on a nutator. Unbound antibodies were washed three times with 1 mL of dig-wash buffer. 100 μl of (1:250 diluted) protein-A-Tn5 loaded with adapters in dig-300 buffer (20 mM HEPES pH 7.5, 300 mM NaCl, 0.5 mM spermidine with Roche cOmplete EDTA-free protease inhibitor) was added to the samples, placed on a nutator for 1 hour and washed three times in 1 ml of dig-300 buffer to remove unbound pA-Tn5. 300 μL tagmentation buffer (Dig-300 buffer + 5 mM MgCl2) was added while gently vortexing, and samples were incubated at 37°C for 1 hr. Tagmentation was stopped by adding 10 μL of 0.5 M EDTA, 3μL of 10% SDS, and 2.5 μL of 20 mg/mL Proteinase K to each sample. Samples were mixed by full speed vortexing for ∼2 seconds and incubated for 1 hr at 55°C to digest proteins. DNA was purified by phenol: chloroform extraction using phase-lock tubes (Quanta Bio) followed by ethanol precipitation. Libraries were prepared using NEBNext HiFi 2x PCR Master mix (M0541S) with a 72°C gap-filling step followed by 13 cycles of PCR with 10-second combined annealing and extension to enrich short DNA fragments. Libraries were sequenced on a Novaseq 6000 (Novogene) with 150 bp paired-end reads.

### Analysis of CUT&Tag and CUT&Run data

For the CUT&Tag and CUT&Run, 150bp pair-end reads were pre-processed using Trimgalore with Fastqc, followed by local aligning to the hg38 genome using bowtie2, with the following pair-end parameters: *–*very-sensitive-local–no-unal–no-mixed–no-discordant–phred33 -I 10 -X 700. Mapping statistics are listed in Extended Data1 (Mapping statistics for NGS data). Mapped bam files were sorted and indexed using samtools ^75^. For peak calling, the bedgraphs were generated using the SEACR pipeline^76^. Bigwig signal files as CPM were generated using the deepTools bamCoverage tool with CPM normalization, along with *–binSize 20 –normalize, using BPM or CPM–scaleFactor = 1.0–smoothLength 60–extendReads 150–centerReads* options. All CUT&Tag or CUT&Run were performed for at least two replicates. For replicates, the data were merged at the BAM level with samtools, followed by a similar processing pipeline to generate bigwigs or bedgraph files. Peaks were called using SEACR with options stringent 0.05 cutoff. CUT&Tag or CUT&Run heatmaps were plotted using the plotHeatmap function in deepTools on the matrix obtained from the computeMatrix output. Peak overlaps for the BRD4 peaks across publicly available ChIP-seq datasets were performed in the Cistrome-DB toolkit browser using the Transcription Factor option^77^. Peaks were overlapped using Intervene across multiple peak sets, including BRD4, E2F6, HDAC2, MAX, MYC, EED, RYBP, H3K14ac, H3K23ac, and H2Aub1, and a correlation heatmap or upset plot was generated using the shiny version of Intervene. The enrichment of these peaks for ChromHMM states in H9 ESC was carried out using the peak2ann function from the genomation package in R. By calculating the overlap of BRD4 peaks on the 15 marks chromHMM states model (hg38 lifted) in H9-ESCs (E008) obtained from the Roadmaps epigenomics project.

### Identification of Active, Bivalent, PRC1.6 target genes

To classify the genes as bivalent or active or PRC1.6 regulated, a heatmap for bivalent marks (H3K7me3 and H3K4me3), PRC1.6 marks (PCGF6, MYC, MAX, E2F6) and active marks (H3K27ac and H3K27ac) was compared along with EED, BRD2, BRD3, BRD4, H3K14ac and H3K23ac modifications in human ESCs. Clustering was performed to identify groups of genes based on the relative abundance of these modifications at the TSSs of protein-coding genes. Similar plotting was performed for bivalent active and H3K27me3-repressed promoters across human ESCs identified (data from ^37^. Likewise, bivalent genes in H9-derived neurons were identified by plotting H3K27me3 (ENCFF285JOX) and H3K4me3 (ENCFF548EWT) signals.

### Site-directed mutagenesis

His_6_-tagged BRD4-BD1 (aa 57-168) and BRD4-BD2 (aa 333-460) constructs in the pNIC28-Bsa4 vector were gifts from Nicola Burgess-Brown (Addgene plasmid #38942 and #38943, respectively). Missense mutations in both bromodomains were generated by site-directed mutagenesis (GenScript) and confirmed by Sanger sequencing.

### RTqPCR

RNA isolation was done using a NEB kit (Monarch, T2040S) followed by reverse transcription using LunaScript RT SuperMix Kit (NEB, E3010), RT–qPCR was done with three independent biological replicates, StepOnePlus Real-Time PCR System (Applied Biosystems). Data were normalized to β-actin (Forward primer CAGCCATGTACGTTGCTATCCAGG, reverse primer AGGTCCAGACGCAGGATGGCATG) from three biological replicates. PRC was performed using EGR1 forward primer CATAGGGAAGCCCCTCTTTC and reverse CTTGTGGTGAGGGGTCACTT, EGR1 forward primer, LMO2 forward primer CACCCCTTACTCCCCATGTT LMO2 reverse primer TTGGCCGCATCTGTTTTCTG

### Protein purification

BRD4-BD1 WT, BRD4-BD1 N140A, BRD4-BD1 D145G, BRD4-BD2 WT, BRD4-BD2 Y390C, BRD4-BD2 Y430C, and BRD4-BD2 N433A were recombinantly purified from BL21(DE3) *E. coli* by nickel-affinity chromatography. BL21(DE3) cells were transformed and grown at 37 °C in 2XYT or Terrific Broth medium with 50 mg/L kanamycin to an optical density of 0.4-0.6 at 600 nm. Bacterial cultures were then grown at 18 °C to an optical density of 0.6-0.8 at 600 nm. Protein expression was then induced overnight with 0.1 mM IPTG at 18 °C. Cells were harvested *via* centrifugation at 5,000 rpm, and cell pellets were frozen at -80 °C until lysis. Frozen cells were thawed on ice and resuspended in lysis buffer (50 mM HEPES, 500 mM NaCl, 5% v/v glycerol, 5 mM imidazole, pH 7.5). 0.3 nM aprotinin, 1 nM E-64, 1 nM leupeptin, 1 nM bestatin, 1 nM pepstatin A, and 100 nM PMSF protease inhibitors were added to the lysis buffer used to resuspend frozen cells, along with 5 µg/mL DNase I and 5 mM MgCl_2_. Resuspended cells were immediately lysed via sonication for 10 minutes (pulsed, amplitude 3.5, 50% work cycle), and lysates were clarified by centrifugation for 30 minutes at 30,000 rpm. Clarified lysates were then applied to nickel-nitrilotriacetic acid (Ni-NTA) resin (∼1 mL resin per L of culture) at 4 °C overnight while rocking. The protein-bound Ni-NTA resin was applied to a column and washed with 15 mL fractions of lysis buffer until nonspecific protein elution had ceased, as determined by monitoring the absorbance of the collected wash fractions at 280 nm. Protein was then eluted from the Ni-NTA resin using increasing concentrations of imidazole in lysis buffer (5 mL of 50, 100, 150, 200, and 250 mM imidazole). Fractions were resolved by SDS-PAGE, and those containing the BRD4 bromodomains of interest were pooled. BRD4-BD1 and -BD2 missense variants were dialyzed overnight in 6,000-8,000 Da molecular weight cutoff dialysis tubing (Fisher Scientific 08-670C) in 4L of storage buffer (50 mM HEPES, 500 mM NaCl, 5% v/v glycerol, pH 7.5), gently stirring at 4°C. BRD4-BD1 and -BD2 WT were further purified by gel filtration using an Enrich SEC 70 10 × 300 mm column (Bio-Rad) or a Superdex 75 Increase 10/300 GL column (Cytiva, 29148721) into storage buffer (50 mM HEPES, 500 mM NaCl, 5% v/v glycerol, pH 7.5). Monomeric protein was collected based on the chromatographs resulting from gel-filtration size-exclusion chromatography. Concentrations of all purified proteins were determined by the Bradford method using bovine serum albumin as a standard, aliquoted, flash-frozen, and stored at -80 °C.

### AlphaScreen assays

All AlphaScreens were conducted in 96-well, ½ Area AlphaPlates (Revvity, 6002350) in a total volume of 40 µL. All stock solutions were prepared in AlphaScreen assay buffer, which consisted of 1x AlphaLISA Epigenetics buffer (Revvity, AL008F), 0.05% v/v Tween 20, and 2 µM tris(2-carboxyethyl)phosphine. Recombinant His_6_-tagged BRD4-BD1 and BRD4-BD2 missense variants were assessed for the ability to bind to a biotinylated histone H3K14ac/23ac(1-27) peptide H_2_N-ARTKQTARKSTGG-K(ac)-APRKQLAT-K(ac)-AGG-K(biotin)-COOH (CPC Scientific, HIST-035A) or a biotinylated H4K5ac/8ac/12ac/16ac (1-18) peptide (Epigentek, R-1008-200). For preliminary screens, 10 µL of a 4× protein stock (final assay concentration 0.07-7 µM for BRD4-BD1 and 0.1-10 µM for BRD4-BD2) was incubated with 10 µL of a 4× peptide stock (final assay concentration 100 nM) in technical triplicate. The plate was incubated at room temperature for 30 minutes. A bead solution containing 8 µg/mL of streptavidin-coated donor beads and 8 µg/mL of nickel-chelate acceptor beads (Revvity, AlphaScreen Histidine (Nickel Chelate) Detection Kit, 500 assay points, 6760619C) was prepared in the assay buffer. Under green-filtered lighting, 20 µL of the bead mixture was added to each well, and the plate was covered and incubated at room temperature for 1 hour. The luminescence of each well was quantified using the Alpha filter cube (Agilent, 1325000) on a BioTek Cytation 5 imaging reader (Agilent, 16277). The resulting Alpha counts were analysed with GraphPad Prism. Alpha counts for initial screens were corrected by subtracting the Alpha counts from the average of the negative control samples (100 nM peptide alone and 10 µM protein alone). Corrected Alpha counts of BRD4-BD1/2 missense variants were then normalised to those of the wild-type bromodomains. For full titrations, 10 µL of a 4× protein stock (final assay concentration 0.001-7 µM for BRD4-BD1 and 0.001-1 µM for BRD4-BD2) was incubated with 10 µL of a 4× peptide stock (final assay concentration 100 nnM) in technical triplicate, and the assay was conducted as above. Raw Alpha counts were transferred to GraphPad Prism and fit to a sigmoidal non-linear regression, then normalized based on the bottom and top values of the fit, to determine the half-maximal effective concentration of binding (EC_50_).

### Public datasets used for ChIP-seq/4su-seq/RNA-seq

#### Public ChIPseq data peaks and bigwigs

Datasets for multiple transcription factors were obtained from public databases, including ENCODE, ChIP-Atlas, and NCBI GEO. This includes peaks for E2F6 (SRX190355, H1), HDAC2 (SRX100390, H1), MAX (SRX190354, H1), MYC (SRX150588, H1), RYBP (SRX3256865, H1), H3K14ac (SRX037086, NCCIT), H3K23ac (SRX17629284, H1) and H2Aub (SRX156117, H1). Same datasets were also used for bigwigs apart from CBX8 (ENCFF189EMD), EZH2 (ENCFF109KCQ), H3K23ac (this study) and H3K14ac (ENCFF079HBG), for which ENCODE bigwigs were used for plotting. For PCGF6, we used the GSE173690 dataset (PGP1 iPSC cell line), where the signal was merged across two replicates. A correlation heatmap of these TFs and histone marks, along with CUT&Tag or CUT&Run peak datasets, was generated using Intervene package using pairwise intersection followed by plotting correlation heatmap showing Kendall’s correlation coefficient, ordered according to PCA using shiny-intervene. Comparison of BRD4 CUT&Tag data at TSS of protein-coding genes and at active enhancers in H9 and H1 from SEDB 2.0 (super enhancer database) was done using deepTools. Other ENCODE datasets ENCFF285JOX (H3K27me3) and ENCFF548EWT (H3K4me3) bigwigs from H9-derived neurons were used for plotting and visualization.

#### Public ChIPseq data reanalysed

For HEK293, the ChIPseq data was reanalysed for PCGF6, MGA, E2F6 and L3MBTL2 downloaded from EBI Array express for accession ID E-MTAB-6006 ^16^. Data was mapped to hg38 genome as described earlier for CUT&Tag data followed by peak calling using MACS3 with default settings using IgG as background with --no model option. For BRD4, peaks and bigwig for HEK293T were downloaded from ChIP-Atlas with accession ID SRX367313. For mESCs, we downloaded PCGF6, RING1B, SUZ12, PCGF1, PCGF2, H3K4me3, H2AUb1, H3K27me3 ChIPseq data on E14 cells with GEO Accession GSE122715. For BRD4 WT and Y430C mutant, we downloaded ChIPseq data from GSE130659. We mapped the data on MM39 using bowtie2 with default parameters for local mapping using single or pair-end mapping option depending on the kind of sequencing. Bam files were merged for the replicates followed by converting them to bed file. Peaks were called using MACS3 with the default parameters with --no model option.

Bigwigs for all ChIP-seq datasets re-analysed were generated using deepTools bamCoverage function with Counts Per Million (CPM) normalisation. For Heatmap, matrices were generated for ChIP-seq peaks in mESCs or HE293/T using computeMatrix function followed by using plotHeatmap function for heatmap. Respective peaks compared were highlighted with red border. Further, TSS’s of genes overlapping PCGF6 peaks in mESCs were obtained using bedtools intersect.

### RNA-seq

For Bulk RNA-seq, fastq files were subjected to QC and trimmed for adapters using Trim-galore, followed by mapping to hg38 reference using STAR-aligner following Bluebee corall pipeline with these parameters: --*peOverlapNbasesMin 40 --peOverlapMMp 0.8 --outFilterType BySJout --outFilterMultimapNmax 200 --alignSJoverhangMin 8 -- alignSJDBoverhangMin 1 --outFilterMismatchNmax 999 --outFilterMismatchNoverLmax 0.6 --alignIntronMin 20 --alignIntronMax 1000000 --limitOutSJcollapsed 5000000 -- seedPerWindowNmax 10.* The counts obtained were then used for the differential expression analyses. For ZxH-treated cells, expression was compared at 4 h, 8 h, and 20 h. The count matrix was used for differential expression, followed by annotation of the DEGs using the integrated Differential Expression and Pathway analysis (iDEP2) tool with default filtration settings. After initial filtering for low coverage and data normalization, differential expression analysis was performed with DESeq2. The different comparisons used for RNA- seq analyses include ZxH (4hr, 8hr, or 20hr) vs DMSO in H9 cells; dTAGV-1 vs H9-WT (DMSO) (20hrs); and BRD4 BD2 mut1 vs WT in H9 cells or H9-derived Neurons. Functional annotation of up- or down-regulated genes was performed using shinyGO or Metascape. For all samples, except the ZxH 4-hour time-point (DMSO has two replicates), have a minimum of 3 replicates. For Gene Set Enrichment Analysis (GSEA), fold-change values were used in the Web-based Gene Set Analysis Toolkit (WEBGESTALT). Genes exhibiting significant deregulation (padj < 0.05) in at least two ZxH treatment time points were filtered and used to compare log2 fold change values as a heatmap using the Morpheus tool for ZxH time points, along with dTAGV-1 vs WT (Extended Data 2, Fig. 1 Heatmap genes). The four clusters obtained were functionally annotated using ShinyGO for GO: Biological Processes, ARCHS4 TF-coexpression, and ARCHS4 Tissues. (Extended Data 2, Metascape_res_genes_Fig.2c). MA-plots were generated for normalised expression against log2 Fold-change, for all comparisons except bulk cerebral organoids, significant (*p*-adj < 0.05) bivalent genes were highlighted in red, other genes in blue, and all non-significant genes in grey. For a paired comparison of changes in expression of the bivalent genes across ZxH treatment time points relative to WT, log2 Fold-change values were visualised in an alluvial plot using gg-alluvial. Genes having a range between ± 0.5 were considered no-change, > 0.5 as upregulated and < -0.5 as down-regulated. We then performed GSEA on these common differentially expressed genes (Extended Data 2; GSEA for genes in Fig. 4b). Paired analysis of genes showing similar directional changes in BRD4 mut1/WT and dTAGV-1/DMSO was performed using an UpSet plot and an XY scatterplot of fold-change values.

For bigwig generation, the individual mapped bam files were sorted and indexed using samtools, followed by bamCoverage (deepTools) with the option normalise using RPKM and a bin size of 20 bp. Further, bigwigs for replicates across the same samples were merged, sorted and indexed with samtools, followed by bigwig generation as described above. To generate normalised bigwig files for BD2 Mut1 vs WT, bigwigCompare was used with a bin size of 20 and skipNAs, with log2 normalisation. These signal files were used to plot or visualise on the genome browser.

### Public RNAseq or 4su-seq Analysis

For hESCs, RNAseq data was downloaded and reanalysed for WT and PCGF6 knock-outs (KO1 and KO2) from GSE173690. Reads were aligned to hg38 following the STAR alignment as described earlier for H9 or H1. Counts were obtained using TE counts followed by differential expression analysis using iDEP2 tool. Similarly, GEO2RNAseq analysis was performed for hESCs between WT and EED-KOs data from GSE92625. Differentially expressed significant genes (padj <= 0.05) were filtered followed by clustering them based on commonality between either ZXH-8hr depletion or BRD4 BD2_mut1, PCGF6-KOs, and EED-KOs using Venn diagram. Significantly common upregulated or downregulated genes were functionally annotated using Metascape.

For 4su-seq in mESCs, raw data was downloaded from NCBI-GEO for accession ID GSE130659. Reads were mapped to mm39 following the STAR mapping pipeline with the parameters as used for H9 or H1 for pair-end reads. Bam files obtained were merged, sorted and indexed using samtools. Bigwigs were generated using deepTools bamCoverage function. Further, log2ratio transformed bigwig was generated using bigwigCompare function. These bigwigs were used for plotting the genes as heatmap or visualisation in genome browser.

### scRNA-seq

Cellplex oligos (10x Genomics) were used for cell hashing, and scRNA-seq libraries were generated by using the NextGEM 3’ v3.1 kit (10x Genomics). scRNA-seq libraries were sequenced with 150 bp paired-end reads at Novogene using the Novaseq X platform.

### scATACseq

scATACseq libraries were generated using 10X Single Cell ATAC v2 reagents and sequenced with 50 bp paired-end reads, yielding 50 million reads per library at Novogene using the Illumina NextSeq platform.

### scRNA-seq data analysis

scRNAseq data were mapped using CellRanger and reference GEX genome GRCh38 (2020-A) from 10x Genomics. The filtered matrix was filtered with DoubletFinder ^79^, assuming a 3.1% doublet formation rate and for nFeature_RNA>500 & percent.mt<10. After filtering, we had WT 3641 cells at d41 and 2541 at d63; mut2 3815 cells at d41 and 3899 at d63; and mut3 4293 at d41 and 4097 at d63. Data was then processed with Seurat 5.3.0 ^80^. Data from UNOs at d41 and d63 were processed separately (in the same way as the merged data), then merged using *merge*, normalized using *NormalizeData and FindVariableFeatures,* and scaled with *ScaleData* using all genes and no regression ^48^. To remove batch effects, we used CSS (cluster similarity spectrum) ^46^. To do so, *cluster_sim_spectrum* was used with the argument *label_tag=” TimePoint”. RunUMAP* was then run on the CSS reduction, with the dims argument set to *1:ncol(Embeddings(obj,” css*”)), as described in the CSS vignette. Finally, the data were clustered first with *FindClusters* at a resolution of 0.2, and the clusters were manually annotated using markers from ^48,49,81^.

### Mapping of scATACseq

scATACseq data were mapped using Cellranger-ATAC and the reference ARC genome GRCh38 (2020-A-2.0.0) from 10x Genomics. Data were then processed with ArchR ^82^ and following the ArchR manual (https://www.archrproject.com/bookdown/index.html). Briefly, arrows were created using minTSS=4 and minFrags=1000. Doublets were identified using *addDoubletScores* and *k =10*. After filtering, we had 4236 cells for WT and 4103 cells for BRD4 mut2. scRNA-seq data were integrated into scATAC-seq using *addGeneIntegrationMatrix*. Differentially accessible peaks were obtained using *addReproduciblePeak* with Macs2 and *getMarkerFeatures*, with log10(nfrags) as the bias, between WT and BRD4mut2. Bigwig files were generated using *getGroupBW* with a tile size of 20 and normalised by the number of cells (nCells).

### Motif enrichment analysis of scATAC-seq data

Normalised ATAC-seq bigwigs for BRD4-BD2 mutant/WT were generated using the deepTools compareBigwig function for each cluster. We filtered for differentially accessible (DA) open peaks in mutants within each cluster, applying a p-value threshold of < 0.01 and a log2FoldChange threshold of > 0.5. For motif enrichment analysis, we used the gimme maelstrom tool to identify differentially enriched motifs within each cluster for the open peaks. We performed central motif enrichment analysis using the Centrimo tool (MEME suite) on the DA open peaks in the BRD4 BD2 mutant for each cell cluster. Data plotted for Gimme maelstrom and Centrimo are listed in Source Data 3.

### scCUT&Tag data

scCUT&Tag data from Zenk et al were used to determine which genes were bivalently marked in UNO populations ^49^. We decided to use the merge dataset from d35, d60, and d120, and the BigWig files for H3K4me3, H3K27ac, and H3K27me3 at those time points, which were generated using *getGroupBw* from archR with a tile size of 100 and normalised by reads in TSS (ReadsInTSS).

### Reporting summary

Further information on research design is available in the Nature Portfolio Reporting Summary linked to this article.

## Data availability

Data is submitted to NCBI GEO and the Broad single-cell portals. Available with the following accession numbers: CUT&Tag and CUT&Run data; GSE313479. Reviewer access: **qxuhaoashfgnrgn**.

Bulk RNAseq; GSE313810, Reviewer access: obonycsqzrqfdev scRNAseq; **SCP3425**

scRNAseq:URL: https://singlecell.broadinstitute.org/single_cell/reviewer_access/54079f88-4f30-4036-b567-0d71264bb526 PIN: HMIYUTJGBU

scATACseq; GSE314451 Reviewer access: ozcxuugafburfev

All the datasets generated and public datasets used in this study are detailed in Extended Table 1. Source data are provided with this paper.

## Code availability

All the analyses in this manuscript have been carried out using publicly available tools. No custom code was generated for this purpose. The methodology contains the details of the analysis steps involved.

## Acknowledgements

We thank Wendy Bickmore and Rob Illingworth (University of Edinburgh), Elena Torlai Triglia, Mirjana Efremova and other members of the QMUL epigenetics centre for their discussion and feedback. We thank Sara Badodi, Radu Zabet, Hemanth Tummala (QMUL), Michael-Christopher Keogh (EpiCypher), Silvia Santos (Francis Crick Institute), Haruhiko Koseki (Riken, Japan), Justyna McIntyre (Polish Academy of Sciences, Poland) and Wei Jiang (Nanchang University, China) for sharing reagents. We thank Xianghua (Cici) Li (King’s College London) and Jeyaprakash Arulanandam (University of Edinburgh) for help in AlphaFold modelling. We thank the Blizard core facilities, including the Genome Centre, for scRNA-seq and scATAC-seq library preparation. This research utilised Queen Mary’s Apocrita HPC facility, supported by QMUL Research-IT.

## Author contributions

Conceptualisation, PMM, MP, and FB; Methodology, FB, MP, PMM, ZS, NA, and AM; Software, MP and FB; Investigation, FB, MP, PMM, ZSZ, NA, AI, PD, DP, MDL, and KLB; Writing – Original Draft, PMM, FB, and MP; Visualisation, FB, MP, MDL, and PMM; Supervision, PMM, DN, and BCS; Funding Acquisition, BCS and PMM.

## Declaration

The authors declare no conflict of interest.

## References

1. Sanchez, R., Meslamani, J. & Zhou, M. M. The bromodomain: From epigenome reader to druggable target. Biochimica et Biophysica Acta - Gene Regulatory Mechanisms vol. 1839 676–685 Preprint at 10.1016/j.bbagrm.2014.03.011 (2014).

2. Hsu, S. C. et al. The BET Protein BRD2 Cooperates with CTCF to Enforce Transcriptional and Architectural Boundaries. Mol. Cell 66, 102–116.e7 (2017).

3. Daneshvar, K. et al. lncRNA DIGIT and BRD3 protein form phase-separated condensates to regulate endoderm differentiation. Nat. Cell Biol. 22, 1211–1222 (2020).

4. Linares-Saldana, R. et al. BRD4 orchestrates genome folding to promote neural crest differentiation. Nat. Genet. 53, 1480–1492 (2021).

5. Donati, B., Lorenzini, E. & Ciarrocchi, A. BRD4 and Cancer: Going beyond transcriptional regulation. Molecular Cancer vol. 17 Preprint at 10.1186/s12943-018-0915-9 (2018).

6. Shi, J. & Vakoc, C. R. The Mechanisms behind the Therapeutic Activity of BET Bromodomain Inhibition. Molecular Cell vol. 54 728–736 Preprint at 10.1016/j.molcel.2014.05.016 (2014).

7. Lovén, J. et al. Selective inhibition of tumor oncogenes by disruption of super-enhancers. Cell 153, 320–334 (2013).

8. Sabari, B. R. et al. Coactivator condensation at super-enhancers links phase separation and gene control. Science (1979). 361, (2018).

9. Han, X. et al. Roles of the BRD4 short isoform in phase separation and active gene transcription. Nat. Struct. Mol. Biol. 27, 333–341 (2020).

10. Conrad, R. J. et al. The Short Isoform of BRD4 Promotes HIV-1 Latency by Engaging Repressive SWI/SNF Chromatin-Remodeling Complexes. Mol. Cell 67, 1001–1012.e6 (2017).

11. Wu, S. Y. et al. Brd4 links chromatin targeting to HPV transcriptional silencing. Genes Dev. 20, 2383–2396 (2006).

12. Zhao, L. et al. BRD4 PRC2 represses transcription of T helper 2 specific negative regulators during T cell differentiation. EMBO J. 42, (2023).

13. Sakamaki, J. ichi et al. Bromodomain Protein BRD4 Is a Transcriptional Repressor of Autophagy and Lysosomal Function. Mol. Cell 66, 517–532.e9 (2017).

14. Kang, H. et al. Bivalent complexes of PRC1 with orthologs of BRD4 and MOZ/MORF target developmental genes in Drosophila. Genes Dev. 31, 1988– 2002 (2017).

15. J.M. Gahan, F. R. & C. E. S. The genetic basis for PRC1 complex diversity emerged early in animal evolution . Proc. Natl. Acad. Sci. U.S.A. 117, 22880–22889, (2020).

16. Stielow, B., Finkernagel, F., Stiewe, T., Nist, A. & Suske, G. MGA, L3MBTL2 and E2F6 determine genomic binding of the non-canonical Polycomb repressive complex PRC1.6. PLoS Genet. 14, (2018).

17. Bryan, E. et al. Nucleosomal asymmetry shapes histone mark binding and promotes poising at bivalent domains. Mol. Cell 10.1016/j.molcel.2024.12.002 (2024) doi:10.1016/j.molcel.2024.12.002.

18. Fallah, M. S., Szarics, D., Robson, C. M. & Eubanks, J. H. Impaired Regulation of Histone Methylation and Acetylation Underlies Specific Neurodevelopmental Disorders. Frontiers in Genetics vol. 11 Preprint at 10.3389/fgene.2020.613098 (2021).

19. Olley, G. et al. BRD4 interacts with NIPBL and BRD4 is mutated in a Cornelia de Lange-like syndrome. Nat. Genet. 50, 329–332 (2018).

20. Deardorff, M. A. et al. HDAC8 mutations in Cornelia de Lange syndrome affect the cohesin acetylation cycle. Nature 489, 313–317 (2012).

21. Basilicata, M. F. et al. De novo mutations in MSL3 cause an X-linked syndrome marked by impaired histone H4 lysine 16 acetylation. Nat. Genet. 50, 1442– 1451 (2018).

22. Arboleda, V. A. et al. De novo nonsense mutations in KAT6A, a lysine acetyl-transferase gene, cause a syndrome including microcephaly and global developmental delay. Am. J. Hum. Genet. 96, 498–506 (2015).

23. Korb, E., Herre, M., Zucker-Scharff, I., Darnell, R. B. & Allis, C. D. BET protein Brd4 activates transcription in neurons and BET inhibitor Jq1 blocks memory in mice. Nat. Neurosci. 18, 1464–1473 (2015).

24. Olley, G. et al. Cornelia de Lange syndrome-associated mutations cause a DNA damage signalling and repair defect. Nat. Commun. 12, (2021).

25. Nowak, R. P. et al. Plasticity in binding confers selectivity in ligand-induced protein degradation article. Nat. Chem. Biol. 14, 706–714 (2018).

26. Nabet, B. et al. The dTAG system for immediate and target-specific protein degradation. Nat. Chem. Biol. 14, 431–441 (2018).

27. Banaszynski, L. A., Chen, L. chun, Maynard-Smith, L. A., Ooi, A. G. L. & Wandless, T. J. A Rapid, Reversible, and Tunable Method to Regulate Protein Function in Living Cells Using Synthetic Small Molecules. Cell 126, 995–1004 (2006).

28. Azuara, V., et al. Chromatin Signatures of Pluripotent Cell Lines. NATURE CELL BIOLOGY vol. 8 (2006).

29. Bernstein, B. E. et al. A Bivalent Chromatin Structure Marks Key Developmental Genes in Embryonic Stem Cells. Cell 125, 315–326 (2006).

30. Lan, X. et al. PCGF6 controls neuroectoderm specification of human pluripotent stem cells by activating SOX2 expression. Nat. Commun. 13, (2022).

31. Shan, Y. et al. PRC2 specifies ectoderm lineages and maintains pluripotency in primed but not naïve ESCs. Nature Communications 8, (2017).

32. Zijlmans, D. W. et al. PRC1 and PRC2 proximal interactome in mouse embryonic stem cells. Cell Rep. 44, (2025).

33. Arnold, M., Bressin, A., Jasnovidova, O., Meierhofer, D. & Mayer, A. A BRD4-mediated elongation control point primes transcribing RNA polymerase II for 3′-processing and termination. Mol. Cell 81, 3589–3603.e13 (2021).

34. Bressin, A. et al. High-sensitive nascent transcript sequencing reveals BRD4-specific control of widespread enhancer and target gene transcription. Nat. Commun. 14, (2023).

35. Zheng, B. et al. Distinct layers of BRD4-PTEFb reveal bromodomain-independent function in transcriptional regulation. Mol. Cell 10.1016/j.molcel.2023.06.032 (2023) doi:10.1016/j.molcel.2023.06.032.

36. Delmore, J. E. et al. BET bromodomain inhibition as a therapeutic strategy to target c-Myc. Cell 146, 904–917 (2011).

37. Court, F. & Arnaud, P. Oncotarget 4110 Www.Impactjournals.Com/Oncotarget An Annotated List of Bivalent Chromatin Regions in Human ES Cells: A New Tool for Cancer Epigenetic Research. Oncotarget vol. 8 www.impactjournals.com/oncotarget/ (2017).

38. Pal, D. et al. H4K16ac activates the transcription of transposable elements and contributes to their cis-regulatory function. Nat. Struct. Mol. Biol. 30, 935–947 (2023).

39. Pradeepa, M. M. et al. Histone H3 globular domain acetylation identifies a new class of enhancers. Nat. Genet. 48, 681–686 (2016).

40. Taylor, G. C. A., Eskeland, R., Hekimoglu-Balkan, B., Pradeepa, M. M. & Bickmore, W. A. H4K16 acetylation marks active genes and enhancers of embryonic stem cells, but does not alter chromatin compaction. Genome Res. 23, 2053–2065 (2013).

41. Lancaster, M. A. & Knoblich, J. A. Generation of cerebral organoids from human pluripotent stem cells. Nat. Protoc. 9, 2329–2340 (2014).

42. Lancaster, M. A. et al. Cerebral organoids model human brain development and microcephaly. Nature 501, 373–379 (2013).

43. Lindenhofer, D. et al. Cerebral organoids display dynamic clonal growth and tunable tissue replenishment. Nat. Cell Biol. 26, 710–718 (2024).

44. Bahrami, S. & Drabløs, F. Gene regulation in the immediate-early response process. Advances in Biological Regulation vol. 62 37–49 Preprint at 10.1016/j.jbior.2016.05.001 (2016).

45. Lanahan, A. & Worley, P. Immediate-Early Genes and Synaptic Function. NEUROBIOLOGY OF LEARNING AND MEMORY vol. 70 (1998).

46. He, Z., Brazovskaja, A., Ebert, S., Camp, J. G. & Treutlein, B. CSS: cluster similarity spectrum integration of single-cell genomics data. Genome Biol. 21, (2020).

47. Becht, E. et al. Dimensionality reduction for visualizing single-cell data using UMAP. Nat. Biotechnol. 37, 38–47 (2019).

48. Kanton, S., Treutlein, B. & Camp, J. G. Single-cell genomic analysis of human cerebral organoids. in Methods in Cell Biology vol. 159 229–256 (Academic Press Inc., 2020).

49. Zenk, F. et al. Single-cell epigenomic reconstruction of developmental trajectories from pluripotency in human neural organoid systems. Nat. Neurosci. 27, 1376–1386 (2024).

50. Fursova, N. A. et al. Synergy between Variant PRC1 Complexes Defines Polycomb-Mediated Gene Repression. Mol. Cell 74, 1020–1036.e8 (2019).

51. Scelfo, A. et al. Functional Landscape of PCGF Proteins Reveals Both RING1A/B-Dependent-and RING1A/B-Independent-Specific Activities. Mol. Cell 74, 1037–1052.e7 (2019).

52. Urli, T. & Greenberg, M. V. C. Epigenetic relay: Polycomb-directed DNA methylation in mammalian development. PLoS genetics vol. 21 e1011854 Preprint at 10.1371/journal.pgen.1011854 (2025).

53. Karmodiya, K., Krebs, A. R., Oulad-Abdelghani, M., Kimura, H. & Tora, L. H3K9 and H3K14 acetylation co-occur at many gene regulatory elements, while H3K14ac marks a subset of inactive inducible promoters in mouse embryonic stem cells. BMC Genomics 13, (2012).

54. Jouret, G. et al. Understanding the new BRD4-related syndrome: Clinical and genomic delineation with an international cohort study. Clin. Genet. 102, 117– 122 (2022).

55. Li, J. et al. BET bromodomain inhibition promotes neurogenesis while inhibiting gliogenesis in neural progenitor cells. Stem Cell Res. 17, 212–221 (2016).

56. Quinonez, S. C. & Innis, J. W. Human HOX gene disorders. Molecular Genetics and Metabolism vol. 111 4–15 Preprint at 10.1016/j.ymgme.2013.10.012 (2014).

57. Blank, M. C. et al. Multiple developmental programs are altered by loss of Zic1 and Zic4 to cause Dandy-Walker malformation cerebellar pathogenesis. Development 138, 1207–1216 (2011).

58. Angelozzi, M. & Lefebvre, V. SOXopathies: Growing Family of Developmental Disorders Due to SOX Mutations. Trends in Genetics vol. 35 658–671 Preprint at 10.1016/j.tig.2019.06.003 (2019).

59. Kikkawa, T. et al. The role of Pax6 in brain development and its impact on pathogenesis of autism spectrum disorder. Brain Research vol. 1705 95–103 Preprint at 10.1016/j.brainres.2018.02.041 (2019).

60. Shoichet, S. A. et al. Haploinsufficiency of novel FOXG1B variants in a patient with severe mental retardation, brain malformations and microcephaly. Hum. Genet. 117, 536–544 (2005).

61. Huber, P. B., Rao, A. & LaBonne, C. BET activity plays an essential role in control of stem cell attributes in Xenopus. Development (Cambridge) 151, (2024).

62. Jin, H. S. et al. Identification of a novel mutation in brd4 that causes autosomal dominant syndromic congenital cataracts associated with other neuro-skeletal anomalies. PLoS One 12, (2017).

63. Groza, T. et al. The International Mouse Phenotyping Consortium: comprehensive knockout phenotyping underpinning the study of human disease. Nucleic Acids Res. 51, D1038–D1045 (2023).

64. Doyle, L. A. et al. RINGs, DUBs and Abnormal Brain Growth—Histone H2A Ubiquitination in Brain Development and Disease. Epigenomes vol. 6 Preprint at 10.3390/epigenomes6040042 (2022).

65. Borges, R. L. et al. Unbalanced chromatin binding of Polycomb complexes drives neurodevelopmental disorders. Mol. Cell 10.1016/j.molcel.2026.01.023 (2026) doi:10.1016/j.molcel.2026.01.023.

66. Gong, F. et al. Screen identifies bromodomain protein ZMYND8 in chromatin recognition of transcription-associated DNA damage that promotes homologous recombination. Genes Dev. 29, 197–211 (2015).

67. Bracken, A. P., et al. EZH2 Is Downstream of the PRB-E2F Pathway, Essential for Proliferation and Amplified in Cancer. The EMBO Journal vol. 22 (2003).

68. Clijsters, L. et al. Cyclin F Controls Cell-Cycle Transcriptional Outputs by Directing the Degradation of the Three Activator E2Fs. Mol. Cell 74, 1264–1277.e7 (2019).

69. Trojer, P. et al. L3MBTL1, a Histone-Methylation-Dependent Chromatin Lock. Cell 129, 915–928 (2007).

70. Mitsuhiro Endoh. PCGF6-PRC1 suppresses premature differentiation of mouse embryonic stem cells by regulating germ cell-related genes. 6:e21064., (2017).

71. Wu, S. Y. et al. Opposing Functions of BRD4 Isoforms in Breast Cancer. Mol. Cell 78, 1114–1132.e10 (2020).

72. Meers, M. P., Bryson, T. D., Henikoff, J. G. & Henikoff, S. Improved cut&run chromatin profiling tools. Elife 8, (2019).

73. Kaya-Okur, H. S. et al. CUT&Tag for efficient epigenomic profiling of small samples and single cells. Nat. Commun. 10, (2019).

74. Langmead, B. & Salzberg, S. L. Fast gapped-read alignment with Bowtie 2. Nat. Methods 9, 357–359 (2012).

75. Danecek, P. et al. Twelve years of SAMtools and BCFtools. Gigascience 10, (2021).

76. Meers, M. P., Tenenbaum, D. & Henikoff, S. Peak calling by Sparse Enrichment Analysis for CUT&RUN chromatin profiling. Epigenetics Chromatin 12, (2019).

77. Zheng, R. et al. Cistrome Data Browser: Expanded datasets and new tools for gene regulatory analysis. Nucleic Acids Res. 47, D729–D735 (2019).

78. Dobin, A. et al. STAR: Ultrafast universal RNA-seq aligner. Bioinformatics 29, 15–21 (2013).

79. Zheng, G. X. Y. et al. Massively parallel digital transcriptional profiling of single cells. Nat. Commun. 8, (2017).

80. Hao, Y. et al. Dictionary learning for integrative, multimodal and scalable single-cell analysis. Nat. Biotechnol. 42, 293–304 (2024).

81. La Manno, G. et al. RNA velocity of single cells. Nature 560, 494–498 (2018).

82. Granja, J. M. et al. ArchR is a scalable software package for integrative single-cell chromatin accessibility analysis. Nat. Genet. 53, 403–411 (2021).

