## Supplementary file for "BRD4 represses developmental and neuronal genes through interaction with PRC1.6" (from another version of this preprint: version 3, 2026-07-30)

**Included Extended data Fig 1-7**

**Extended data Table 1:** Information regarding the NGS datasets used in this paper either generated in this paper or from public databases

**Source Data 1**
**Key Resource Table:** Information regarding the Reagents and materials as well as the software and tools used in the study.
**Mapping statistics for NGS data:** Mapping details of the NGS datasets generated in this study
**Source data for PLA:** Quantification of PLA images foci per cell for the Extended Data Figure 4g

**Source Data 2**
**Fig.1 Heatmap genes:** Genes compared in Figure 1d
**Metascape_res_genes_Fig2c:** Full GO annotation of genes compared in Figure 2c
**Metascape_res_genes_Fig5b&c:** Full GO annotation of genes compared in Figure 5b&c
**Metascape_res_genes_Fig5d:** Full GO annotation of genes compared in Figure 5d

**Source Data 3:** Data for the central motif enrichment analysis (Centrimo) for the TFs shown in Extended Data Figure 7c and GIMME maelstrom analysis of TFs shown in Figure 7b
**Centrimo results RPE**
**Centrimo results Glutamatergic**
**Centrimo results GABAergic2**
**Centrimo results GABAergic**
**Centrimo results G2M**
**Centrimo results Diencephalic2**
**Centrimo results Diencephalic**
**Gimme Motif Enrichment**

a

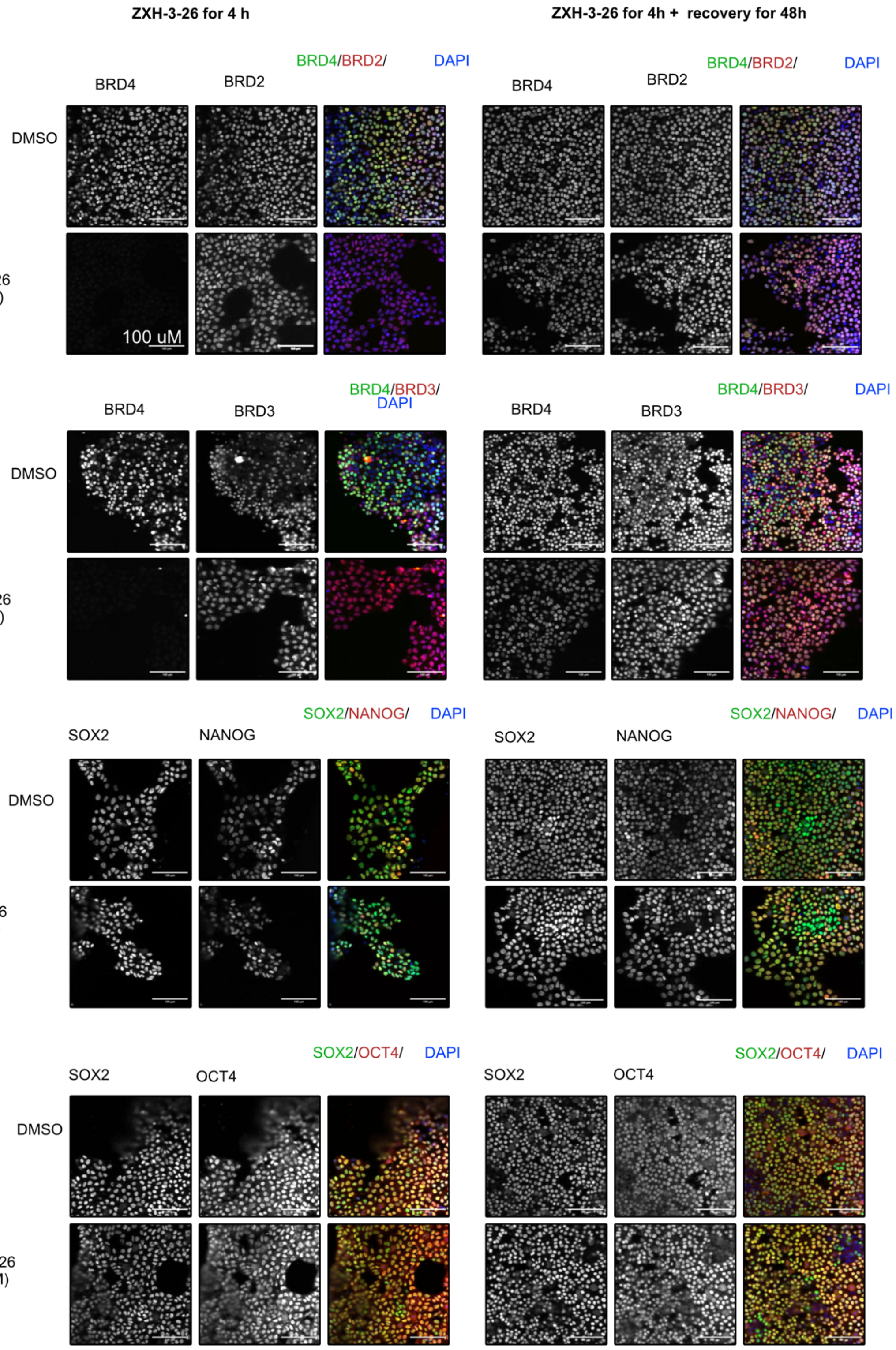

**Extended Data Figure 1: Related to Figure 1**; **a**) Immunofluorescence images showing expression level of pluripotency markers (SOX2 and OCT4) upon DMSO or ZxH-3-26 (ZxH) treatment for 4 hours (top two) and upon rescue or recovery by changing to media without DMSO or ZxH for 48 hours, scale bars: 10 µm.

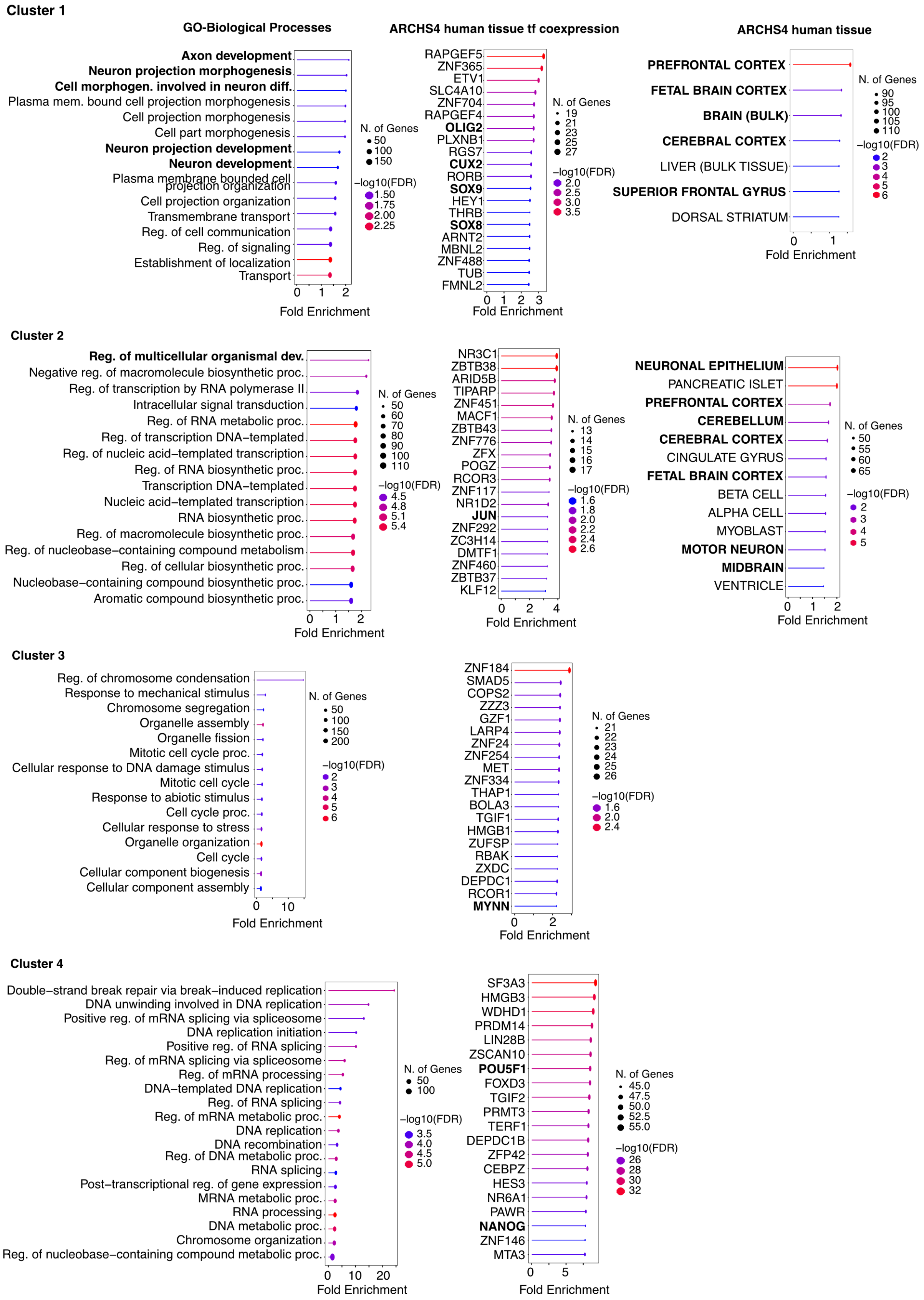

**Extended Data Figure 2: related to Fig. 1d**, enrichment of Gene Ontology (GO) biological processes (left), all RNA-seq and ChIP-seq Sample and Signature Search 4 (ARCHS4) human tissue TF coexpression (middle) and ARCHS4 human tissue enrichment of the genes in the four clusters in Fig. 1d.

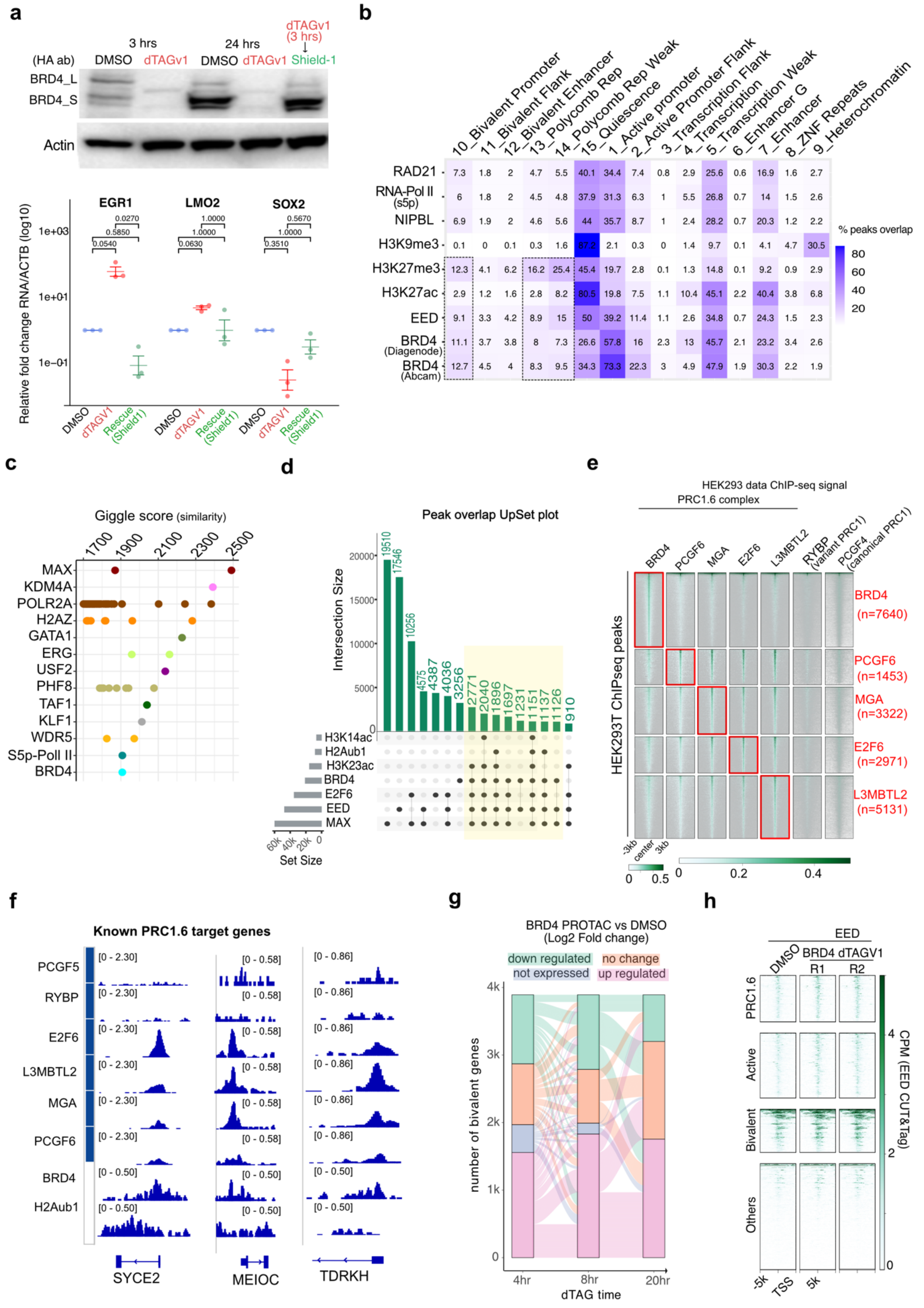

**Extended Data Figure 3: Related to Fig. 1-3**; **a**) Western blots for HA (detects long and short BRD4 isoforms in HA-FKBP-BRD4) hESCs, and ACTIN antibodies for DMSO, dTAGv1 (500uM) treated for 3 hrs followed by degradation rescue by addition of Shield1 for 24 hrs (top). RT-qPCR for EGR1, LMO2 and SOX2 genes in DMSO (3hr), BRD-dTAGV1 (3hrs) and Sheild1 for 24hrs. p-values for paired t-test with Bonferroni correction (bottom). **b**) Percentage peak overlap for BRD4, EED, RAD21, NIPBL, serine-5 phosphorylated RNA Pol II (RNA-Pol II s5p), H3K27ac, H3K4me3, and H3K27me3 across 15 ChromHMM states in H9-hESCs. **c**) Dot-plot showing peak similarity for BRD4 CUT&Tag peaks (as GIGGLE score) across different TF-ChIP-seq peaks for different cell lines (each dot is a cell line) using Cistrome-DB Toolkit. **d**) UpSet plot illustrating overlap of CUT&Run (EED) or CUT&Tag (BRD4) or ChIP-seq peaks (E2F6, MAX, H3K23ac, H3K14ac, H2Aub1). The bar plot at the top shows the size of each intersection, while the bar plot on the left indicates the size of each set. Peaks overlapping between BRD4, PCGF6, EED, H3K4me3, and H3K27me3 are highlighted in yellow. **e**) Heatmaps showing enrichment of BRD4 (HEK293T) and PRC1.6 components ChIP-seq signal for each ChIP-seq peak from HEK293 cells. BRD4 ChIPseq data is from (ChIP-Atlas) and PRC1 components is from Stielow et al 2018 [1]. **f**) Like (e) but genome-browser tracks for BRD4 and PRC1.6 components along with PCGF5 and RYBP ChIP-seq signal at known PRC1.6 target genes. **g**) Alluvial plot showing RNA-seq data for up- and down-regulated ESC bivalently marked genes upon ZxH treatment at 4, 8 and 20 hours. **h)** Heatmap for EED CUT&Tag signal comparison at the genes in DMSO and upon BRD4 depletion for 3hrs (dTAGV1)

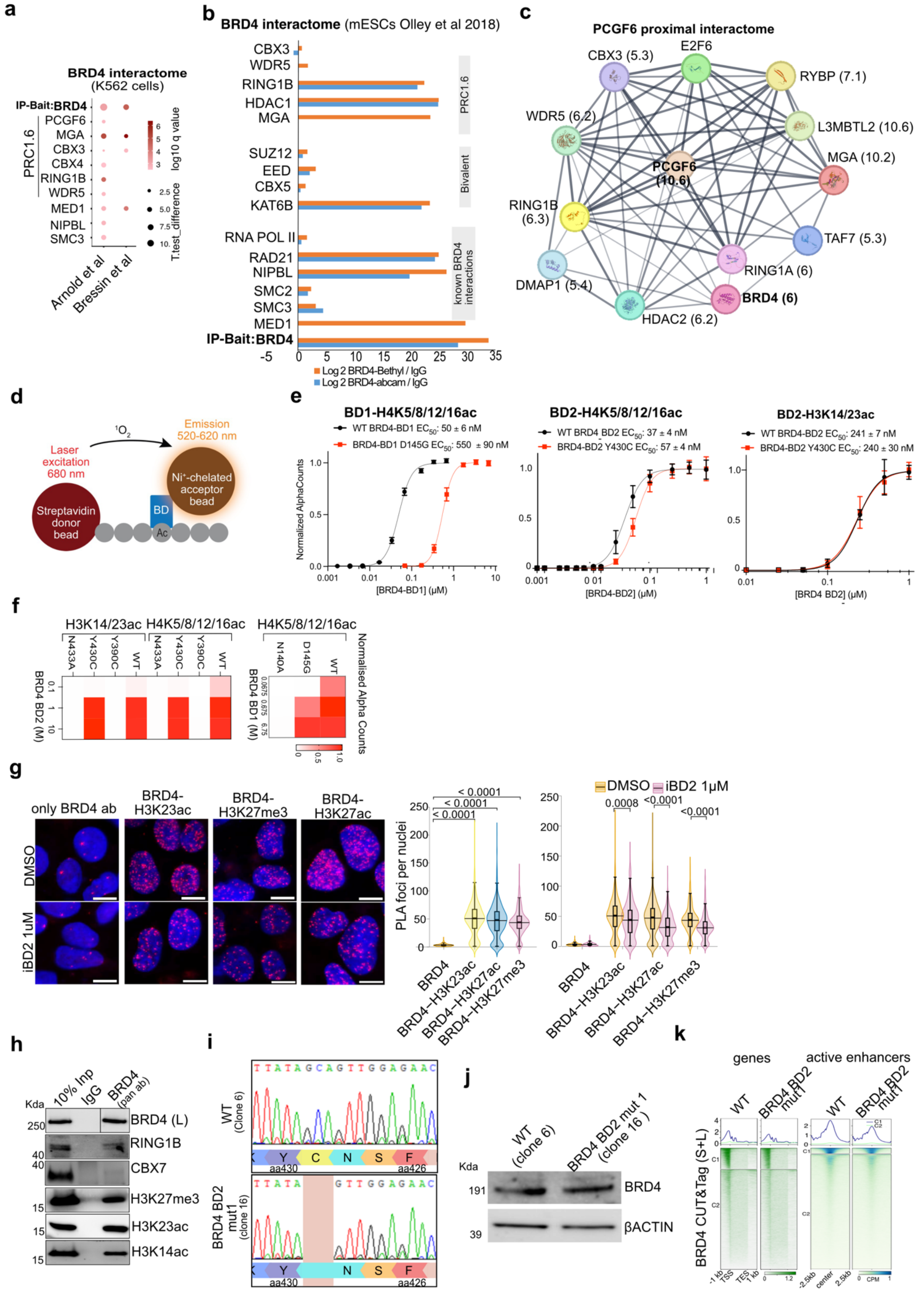

**Extended Data Figure 4: Related to Figs. 3-5; a**) Log2 fold enrichment in BRD4 IP in mESCs over IgG control for PRC proteins, along with known BRD4 interactors, the bar graph shows log2 enrichment of the indicated proteins relative to IgG control in BRD4 IP using independent antibodies (Abcam and Bethyl lab), data from (Olley et al 2018)[2]. **b**) Similar to Fig. 3a but for enrichment of PRC proteins in BRD4 immunoprecipitation from K562 cells, data from (Brassin et al 2023 and Arnold et al 2021) [3,4]. The size of the circle represents the t-test statistic for the difference between the BRD4 IP and the IgG control. c) **c**) STRING map of PRC1.6 proximal interactome (data from PCGF6 Turbo-ID, from Zijlmans et al, 2025) [5] , log2 fold enrichment of each protein in brackets. Line thickness indicates the degree of confidence in predicting the interactions (related to Fig. 3a). **d**) Illustration of the AlphaScreen assay used to measure affinity between BRD4 bromodomains to acetylated histone peptides. **e**) Alpha Screen binding curve showing BRD4 binding affinity to acetylated histone peptides. Binding curves for wild-type (WT) BRD4-BD1 and BRD4-BD2 with H4K5ac/8ac/12ac/16ac and H3K14ac/23ac peptides, comparing $EC_{50}$ values for WT and mutant variants (D145G for BD1, Y430C for BD2). **f**) Heatmap showing normalised Alpha counts for WT and BD1 (D145G) and BD2 (Y430C, N433A) CdLS missense variants with H4K5ac/8ac/12ac/16ac and H3K14ac/23ac peptides, mutation in the essential BC loop of BD1 (N140A) and BD2 (N433A) served as positive control. **g**) Representative proximity ligation assay (PLA) images for α-BRD4 with α-H3K27ac, and α-H3K27me3 antibodies in hESCs with 1μM of inhibitor of bromodomain 2 (iBET-BD2) and DMSO (left), scale bars: 10 μm. The number of PLA foci per cell was quantified and plotted (right, $n > 372$ cells over three independent experiments; horizontal lane is median value that has been indicated, the box spans the interquartile range**,** from the 25th percentile (Q1) to the 75th percentile (Q3); *p*-values obtained from a two-tailed Mann–Whitney test). **h**) Like Fig. 3b, but IP using BRD4 antibody. **i**) Sanger sequencing trace showing deletion of amino acid 429 in BRD4 BD2 mut 1 clone (clone 16), and corresponding wild type clone (clone 6). **j**) Western blot showing BRD4 expression in WT and BRD4 BD2 mut 1 clone, β-ACTIN served as loading control. **k**) Heatmaps showing enrichment of BRD4 ChIP-seq signal (CPM), in WT and BRD4 BD2 mut 1 hESCs at protein coding genes and active enhancers.

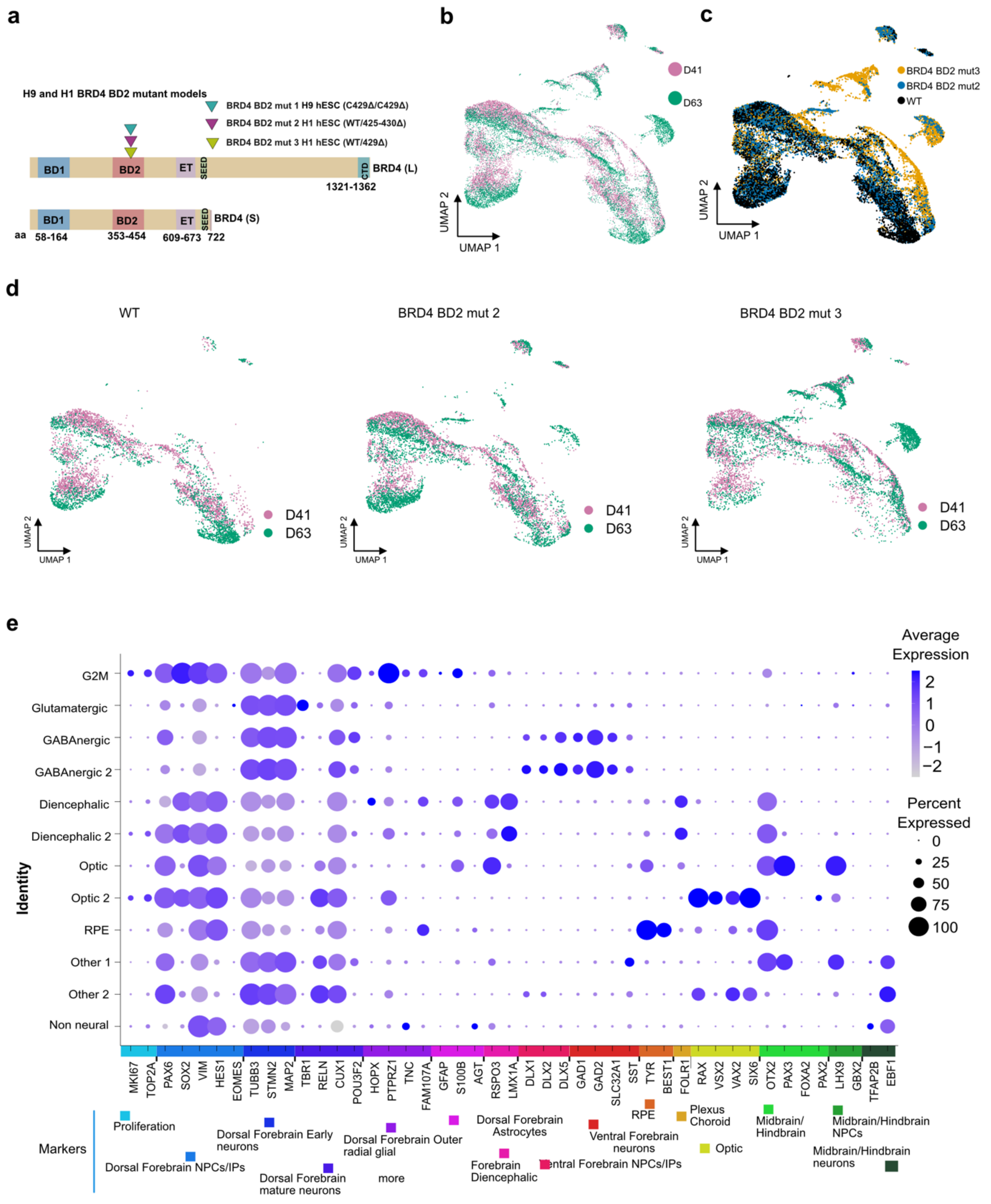

**Extended Data Figure 5**: **Related to Fig. 6**, **a**) Illustration showing the BRD4 domains and homozygous and heterozygous mutations in BD2 of BRD4 in H9 and H1 hESCs. **b**) UMAP visualisation of integrated single-cell RNA sequencing (scRNA-seq) data, showing clustering by 41-day and 63-day unguided neural organoids (UNOs). **c)** UMAP visualisation showing the distribution of cells across genotypes: wild-type (WT), *BRD4* BD2 mutant 2 and *BRD4* BD2 mutant 3. d) Similar to (b) but including time points (day 41 and day 63) for wild-type (WT), *BRD4* BD2 mutant 2, and *BRD4* BD2 mutant 3 in separate UMAPs. **e**) Dot plot displaying average gene expression and percentage of cells expressing key marker genes for various identified cell identities.

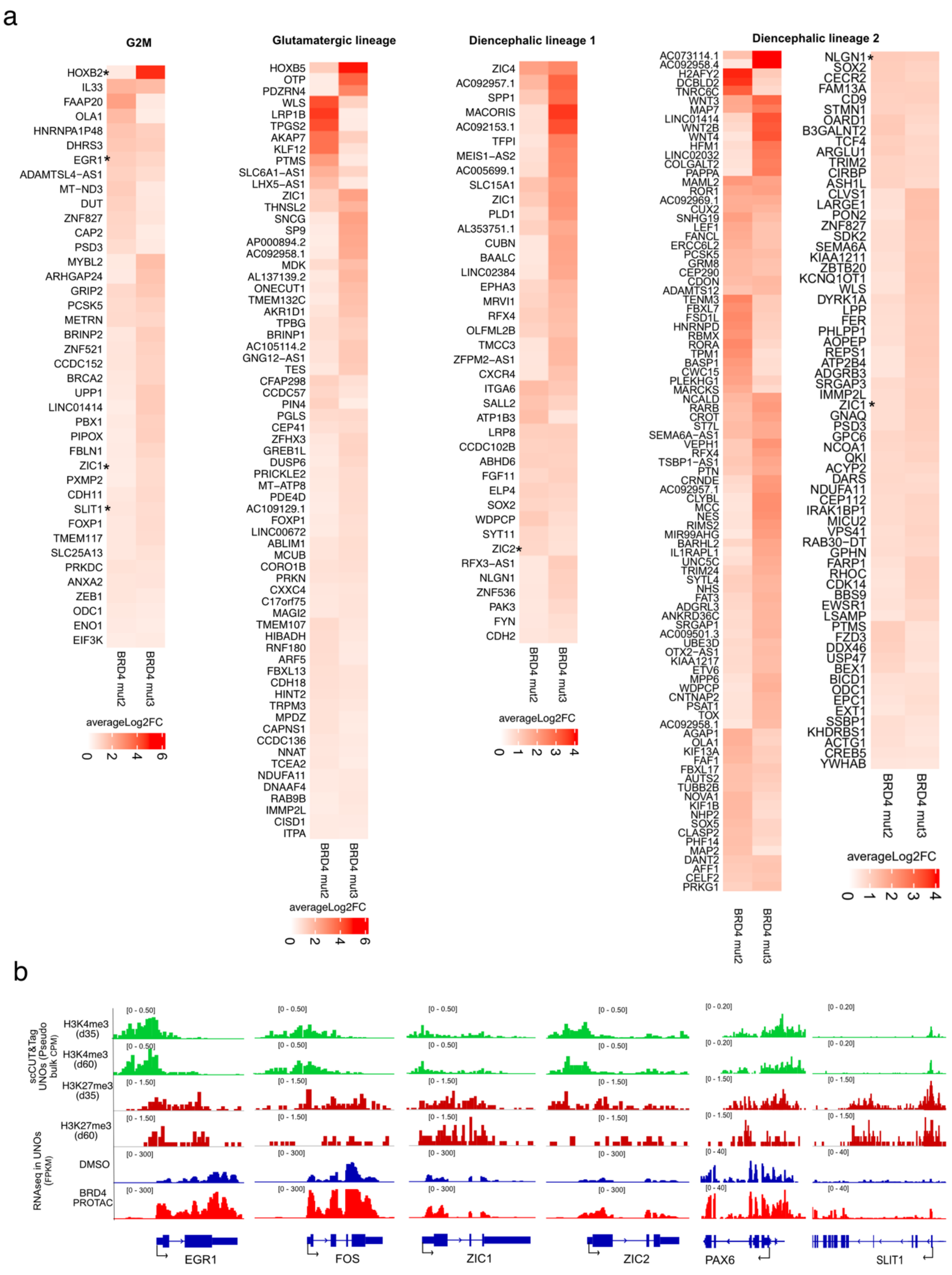

**Extended Data Figure 6: Related to Fig. 6.** **a**) Heatmap showing significantly upregulated genes in scRNAseq data from WT and BRD4 mutant UNOs. Genes significantly upregulated in both mutant (BRD4 BD2 mut 2 and BRD4 BD2 mut 3) and having a log2FC>0.5 or log2FC< -0.5 for Glutamatergic lineage G2M and (d) are shown. **b**) Genome-browser visualisation of pseudo bulk single cell CUT&Tag (scCUT&Tag) counts per million (CPM) for H3K4me3 and H3K27me3 from day 35 and day 60 UNOs, along with average RNAseq signal (FPKM, n=3 replicates), performed 20 hours after DMSO and ZxH treatment in wild type day 41 UNOs at genes that are bivalently marked and upregulated in both BRD4 BD2 mutant UNOs.

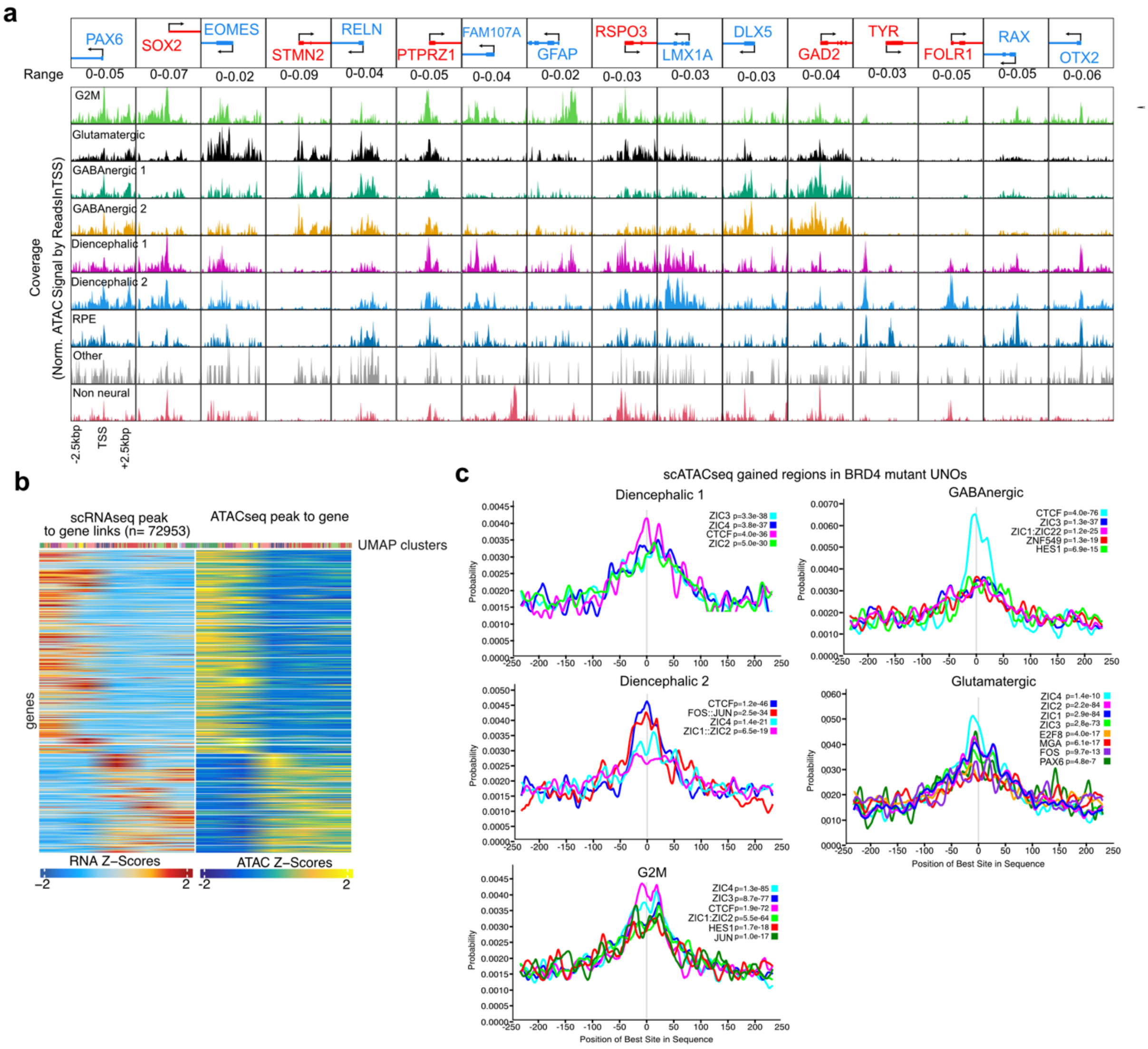

**Extended Data Fig. 7: Related to Fig. 7, a)** Single cell ATAC-seq (scATAC-seq) signal coverage across key marker gene promoters. **b**) Peak to gene heatmap (Z-scores) for scATAC-seq and scRNA-seq data. **c**) c) Meta-plots showing representative motifs centrally enriched in each cell type cluster for the scATAC-seq peaks gained in the BRD4 BD2 mutant, identified using the Centrimo tool. The Y-axis shows the probability of the motifs being enriched at the position on the X-axis, with motifs shown alongside their p-values for each cluster.

### Extended Data Table 1

| **Datasets generated in this paper** | | |
|---|---|---|
| **TF or histone modification or RNAseq** | **Accession ID** | **Source** |
| RNAseq | GSE313810 | This Paper |
| CUT&Tag and CUT&Run seq | GSE313479 | This Paper |
| scRNAseq | scRNAseq:URL: https://singlecell.broadinstitute.org/single_cell/reviewer_access/54079f88-4f30-4036-b567-0d71264bb526 PIN: HMIYUTJGBU | This Paper |
| scATACseq | GSE314451 | This Paper |
| **Public Datasets used**<br>**Peaks and/or bigwigs used in Figure 2a peak correlation heatmap and 2b bigwigs heatmap at protein coding genes** | | |
| E2F6 (Peaks and bigwig) | SRX190355 | |
| HDAC2 (Peaks) | SRX100390 | |
| MAX (Peaks) | SRX190354 | |
| MYC (Peaks and bigwig) | SRX150588 | |
| RYBP (Peaks and bigwig) | SRX3256865 | |
| H3K14ac (Peaks) | SRX037086 | |
| H3K23ac (Peaks) | SRX17629284 | |
| H2A119Ub (Peaks) | SRX156117 | |
| BCOR_H1_peaks_.05.bed | SRX10382441 | 6 |
| JARID2_hPSC_peaks_.05.bed | SRX8171699 | |
| SMC3_peaks_H9.bed | SRX5730049 | |
| CBX3_H1_peaks_.05.bed | SRX3010018 | |
| PCGF2_NT2D1_peaks_.05.bed | SRX16005284 | |
| CBX3_H1_peaks_.05.bed | SRX3010018 | |
| hESC_EED_05 | SRX8175102 | |
| PCGF1_H1.bw | SRX3256864 | |
| KDM2B_H1.bw | SRX3256862 | |
| CBX8 | ENCFF189EMD | ENCODE |
| H1_SUZ12.bw | ENCFF552BOU | ENCODE |
| EZH2 (Peaks and bigwig) | ENCFF109KCQ | ENCODE |
| H3K14ac (Peaks and bigwig) | ENCFF079HBG | ENCODE |
| RNF2_H1_peaks.bed | ENCFF283MNG | ENCODE |
| HEK293T BRD4 ChIPseq peaks and bigwig | SRX367313 | 6 |
| HEK293 PRC1.6 ChIPseq data for PCGF6, MGA and L3MBTL2 | E-MTAB-6006 | 1 |
| **RNAseq data used form NCBI GEO for figure 2c** | | |
| PCGF6 WT, KO-RNAseq counts and bigwigs for PCGF6 | GSE173690 | 7 |
| EED knockout RNAseq data on H1 hESCs | GSE92625 GEO2RNAseq DEseq | 8 |
| **RNAseq data and ChIPseq data for mESCs used form NCBI GEO for figure 5a-c** | | |
| PCGF6, RING1b, SUZ12, PCGF1, PCGF2, H3K4me3, H2AUb1, H3K27me3 ChIPseq data on E14 cells | GSE122715 | 9 |
| WT, BRD4-Y430C ChIPseq and RNAseq on mESCs | GSE130659 | 10 |
| H3K27me3, H3K4me3 H9 | GSE200768 | 11 |
| H3K27me3 Neuron | ENCFF285JOX | ENCODE |
| H3K4me3 Neuron | ENCFF548EWT | ENCODE |

**Supplementary references**

1. Stielow, B., Finkernagel, F., Stiewe, T., Nist, A. & Suske, G. MGA, L3MBTL2 and E2F6 determine genomic binding of the non-canonical Polycomb repressive complex PRC1.6. *PLoS Genet.* **14**, (2018).
2. Olley, G. *et al.* BRD4 interacts with NIPBL and BRD4 is mutated in a Cornelia de Lange-like syndrome. *Nat. Genet.* **50**, 329–332 (2018).
3. Bressin, A. *et al.* High-sensitive nascent transcript sequencing reveals BRD4-specific control of widespread enhancer and target gene transcription. *Nat. Commun.* **14**, (2023).
4. Arnold, M., Bressin, A., Jasnovidova, O., Meierhofer, D. & Mayer, A. A BRD4-mediated elongation control point primes transcribing RNA polymerase II for 3′-processing and termination. *Mol. Cell* **81**, 3589-3603.e13 (2021).
5. Zijlmans, D. W. *et al.* PRC1 and PRC2 proximal interactome in mouse embryonic stem cells. *Cell Rep.* **44**, (2025).
6. Zou, Z., Ohta, T. & Oki, S. ChIP-Atlas 3.0: A data-mining suite to explore chromosome architecture together with large-scale regulome data. *Nucleic Acids Res.* **52**, W45–W53 (2024).
7. Lan, X. *et al.* PCGF6 controls neuroectoderm specification of human pluripotent stem cells by activating SOX2 expression. *Nat. Commun.* **13**, (2022).
8. Shan, Y. *et al.* PRC2 specifies ectoderm lineages and maintains pluripotency in primed but not naïve ESCs. *Nature Communications* **8**, (2017).
9. Scelfo, A. *et al.* Functional Landscape of PCGF Proteins Reveals Both RING1A/B-Dependent-and RING1A/B-Independent-Specific Activities. *Mol. Cell* **74**, 1037-1052.e7 (2019).
10. Olley, G. *et al.* Cornelia de Lange syndrome-associated mutations cause a DNA damage signalling and repair defect. *Nat. Commun.* **12**, (2021).
11. Pal, D. *et al.* H4K16ac activates the transcription of transposable elements and contributes to their cis-regulatory function. *Nat. Struct. Mol. Biol.* **30**, 935–947 (2023).
